# Single nuclei chromatin accessibility and transcriptomic map of breast tissues of women of diverse genetic ancestry

**DOI:** 10.1101/2023.10.04.560911

**Authors:** Poornima Bhat-Nakshatri, Hongyu Gao, Aditi S. Khatpe, Patrick C. McGuire, Cihat Erdogan, Duojiao Chen, Guanglong Jiang, Felicia New, Rana German, Anna Maria Storniolo, Yunlong Liu, Harikrishna Nakshatri

## Abstract

Single nuclei analysis is allowing robust classification of cell types in an organ that helps to establish relationships between cell-type specific gene expression and chromatin accessibility status of gene regulatory regions. Using breast tissues of 92 healthy donors of various genetic ancestry, we have developed a comprehensive chromatin accessibility and gene expression atlas of human breast tissues. Integrated analysis revealed 10 distinct cell types in the healthy breast, which included three major epithelial cell subtypes (luminal hormone sensing, luminal adaptive secretory precursor, and basal-myoepithelial cells), two endothelial subtypes, two adipocyte subtypes, fibroblasts, T-cells, and macrophages. By integrating gene expression signatures derived from epithelial cell subtypes with spatial transcriptomics, we identify specific gene expression differences between lobular and ductal epithelial cells and age-associated changes in epithelial cell gene expression patterns and signaling networks. Among various cell types, luminal adaptive secretory cells and fibroblasts showed genetic ancestry dependent variability. A subpopulation of luminal adaptive secretory cells with alveolar progenitor (AP) cell state were enriched in Indigenous American (IA) ancestry and fibroblast populations were distinct in African ancestry. ESR1 expression pattern was distinctly different in cells from IA compared to the rest, with a high level of ESR1 expression extending to AP cells and crosstalk between growth factors and Estrogen Receptor signaling being evident in these AP cells. In general, cell subtype-specific gene expression did not uniformly correlate with cell-specific chromatin accessibility, suggesting that transcriptional regulation independent of chromatin accessibility governs cell type-specific gene expression in the breast.

## Introduction

Breast cancer shows genetic ancestry dependent variability in incidence and outcome ^1^. For example, while breast cancer incidence is lower in women of African ancestry, age at diagnosis and outcome are distinctly different compared to women of European ancestry. Breast cancer is diagnosed at an earlier age and is more likely to be triple negative breast cancer subtype in women of African ancestry compared to women of European ancestry ^2,3^. Even after controlling for socioeconomic stress and healthcare access, breast cancer outcome tends to be worse in women of African ancestry ^3^. Indigenous Americans (IA) also experience a lower incidence of breast cancer compared to non-Hispanic White women (74.2 per 100,000 versus 126.1 per 100,000) ^4^. A subpopulation of IA women carries a breast cancer protective allele, particularly against Estrogen Receptor-negative (ER-) breast cancer, on chromosome 6q25 close to estrogen receptor 1 (ESR1) gene ^5^. IA women experience a disproportionately higher level of HER2+ breast cancers ^4^. There is evidence in the literature for genetic ancestry dependent differences in mutational spectrum suggesting the influence of genetic ancestry on genome organization and accessibility to undergo genomic changes ^6^.

As a first step in understanding the complex biology of breast cancers, several groups have utilized single cell technologies to develop single cell atlas of the breast ^7–11^. We reported a single cell atlas of the breast utilizing breast tissues donated for research purpose by women with no clinical history of breast cancer ^11^. We described 23 epithelial cell states; 8 basal-myoepithelial, 3 intermediate basal/luminal progenitor, 8 luminal progenitor (recently renamed as luminal adaptive secretory precursor, LASP cells) and 4 mature luminal (recently renamed as luminal hormone sensing, LHS cells) cell states. Gene expression signatures corresponding to three cell states within LHS cells and one within LASP cells are enriched in the majority of breast cancers, suggesting that these cell states are putative cells-of-origin of breast cancer ^11^. Other studies cited above utilized tissues from reduction mammoplasty or tumor adjacent normal tissues to derive single cell atlas of the breast. For example, Kumar et al described 12 major cell types in the breast with epithelial cells under 11 different states (one basal-myoepithelial, seven LASP, and three LHS cell states) ^7^. Gray et al described six epithelial cell states (hormone sensing-alpha, hormone sensing-beta, alveolar progenitor, basal-luminal hybrid, basal-alpha and basal-beta) and suggested these states are influenced by age, parity and BRCA2 mutation status^8^. They also suggested that age impacts the accumulation of alveolar cells with poor transcriptional lineage fidelity. Murrow et al characterized premenopausal breast tissue at the single cell level to determine coordinated transcriptional programs that alter in response to changing hormonal levels ^9^. Two other single cell studies using reduction mammoplasty samples identified three major epithelial cell types in the breast ^10,12^. However, there remains a lack of information in differences in cell state based on genetic ancestry and relationship between cell state as defined by transcriptome and chromatin accessibility status.

In this study, we utilized breast tissues donated by women with ancestries as follows: 22 Ashkenazi-European ancestry, 10 Asian with a mix of East and central South Asia ancestry, 20 European-non-Ashkenazi, 10 Hispanic-White with predominant European ancestry and 10 Indigenous American ancestry. We performed integrated single nuclei ATAC-seq (snATAC-seq) and RNA-seq (snRNA-seq) analysis. Due to technical difficulties, we were able to perform only single nuclei RNA-seq of 10 African ancestry donors. Additional tissues from European ancestry donors were subjected to only snRNA-seq to allow comparison between African and European ancestry. Gene expression signatures derived from our previous single cell RNA sequencing data ^11^ was applied on to spatial transcriptomics data obtained from tissues of three women who donated breast tissues twice, 10 years apart, to identify genes enriched in lobular epithelial cells compared to ductal epithelial cells and to identify individual level gene expression change with age. Our studies suggest that genetic ancestry impacts the epithelial cell state with Indigenous Americans being enriched for cells with alveolar progenitor state. Further genetic ancestry dependent differences in fibroblasts were noted with fibroblasts from African ancestry clustering differently with distinct cell state and gene expression. These results suggest that a comprehensive analysis of every cell type, not just the epithelial cell type, in an organ is needed to study the impact of genetic ancestry on disease incidence, subtypes and progression.

## Results

### Single nuclei chromatin accessibility and transcriptome mapping of breast tissues from women of diverse genetic ancestry

Figure S1 provides a brief overview of the experimental design and Figure 1a provides genetic ancestry estimates of the donor samples used in the study. Details of genetic ancestry estimation analysis are described in our recent publication ^13^. Table S1 provides details of donors including menopausal status, age, BMI, self-reported race, days since ovulation etc. Average age, BMI and number of childbirths in each subgroup are also provided in this table. The majority of self-reported White donors are enriched for European ancestry, whereas the majority of self-reported Black donors are enriched for African ancestry. Sixty percent of self-reported Hispanic women are enriched for European ancestry. Fifty percent of Asians are enriched for East Asian while other 50% are enriched for Southeast Asian ancestry markers. Indigenous Americans are enriched for “Americana” ancestry markers and Americana ancestry proportion in these donors is similar to Indigenous American ancestry proportions described in Indigenous American breast cancer patients in Peru ^4^. Note that this is the only group to carry significant “Americana” ancestry markers. Ashkenazi Jewish Americans are a mixture of European, Middle Eastern, and African ancestry. We also included six samples from BRCA1 mutation donors and five BRCA2 donors and information on these donors are provided in another study that utilized these samples for single cell RNA-seq ^14^. Table S2 provides details of number of nuclei sequenced in each group and number of genes per nuclei per group. Although multiome assay that combined snATAC-seq and snRNA-seq was attempted with tissues from donors of all genetic ancestry, for unknown reasons, combined multiome assay did not work with tissues from donors of African ancestry. Therefore, only snRNA-seq was performed with the tissues of donors of African ancestry. For better comparison of data from breast tissues of African ancestry, a set of samples from parous and nulliparous women of European ancestry was similarly processed to obtain snRNA-seq.

**Figure 1:**
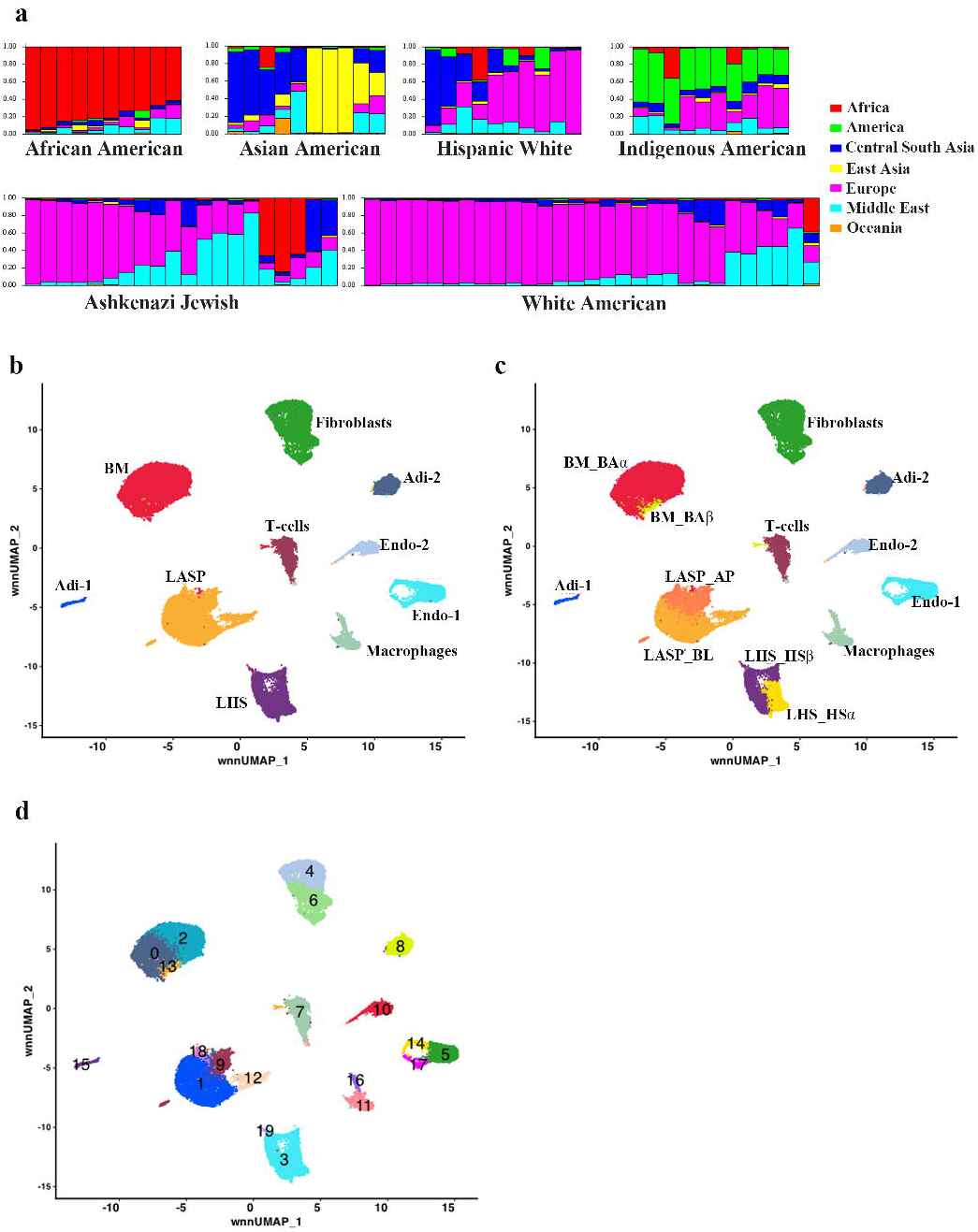
Integrated snATAC-seq and snRNA-seq analyses of breast issues of healthy women. a) Genetic ancestry marker distribution pattern among donors of self-identified race/ethnicity groups. b) Integrated cell clusters generated using snATAC-seq and snRNA-seq data representing all donors except African American donors. c) Breast epithelial cells could be further subclassified into six different cell types. d) Cell clustering analysis reveal further refinement of cell state of LASP cells, fibroblasts, endothelial cells.

Integrated analysis of snATAC-seq and snRNA-seq data from all ancestries except African ancestry revealed 10 distinct clusters: three of them are epithelial cell types, two are endothelial cell types, two adipocyte subtypes, fibroblasts, T cells and macrophages (Figure 1b). Epithelial cell types were annotated using CD49f/EpCAM markers as done previously into mature luminal (ML), luminal progenitors (LP) and basal cells ^11^. These cell clusters have been described with various names in the literature ^7,8,12^. In a recent breast cell annotation event organized by Chan-Zuckerberg Initiative that included several research groups involved in developing single cell atlas of the breast, the following terminologies were suggested: luminal hormone sensing (LHS), luminal adaptive secretory precursor cells (LASP) and basal-myoepithelial (BM) cells for mature luminal, luminal progenitor, and basal cells, respectively. Another recent study that combined single cell RNA-seq with single cell proteome assay has subclassified hormone sensing cells into HS-alpha (HSα) and HS-beta (HSβ), LASP cell into alveolar progenitors (AP) and basal-luminal alveolar (BL) cells and BM cells into Basal-alpha (BAα) and Basal-beta (BAβ) cells ^8^. We used the markers described in that study to subcluster the epithelial cells and found the presence of all six epithelial cell states (Figure 1c). Further examination of these data revealed LASP cells to be in four cell states, fibroblasts in two states, endothelial cells in four states and macrophages in two states (Figure 1d). Genes expressed in each of the cell types are listed in Table S3. Genes differentially expressed in HSα compared to HSβ cells, AP versus BL cells and BAα versus BAβ cells are listed in supplementary Tables S4, S5, and S6, respectively. Genes differentially expressed in different cell states of endothelial cells, fibroblasts and adipocytes are listed in Tables S7, S8, and S9, respectively. The top ten transcription regulators of each of the major cell types are listed in Table S10. Average expression of genes in each cluster is listed in Tables S11.

### AP cells are enriched in breast tissues of women of Indigenous American ancestry

We next analyzed data based on self-reported racial/ethnicity groups. Samples from BRCA1 and BRCA2 mutation carriers were also included. Cell types in each group are shown in Figure 2a and cell proportions are listed in Table S12. Data from women of African ancestry are only from snRNA-seq and additional details for this group are described in subsequent sections. AP cells were disproportionately higher in breast tissues of women of Indigenous American ancestry. While 19% of cells in Indigenous Americans are AP cells, the percentage of AP cells in the other groups ranged from 5 to 9%. Using the previously described markers of alveolar cells (EHF and ELF5) and luminal progenitor cells (KIT) ^15,16^, we confirmed that AP cells express alveolar markers and KIT (Figure 2b-d). None of these differences between the groups is due to differences in age, BMI or number of childbirths as the average age of European, Indigenous American, Asian, and Hispanic-White donors were 38, 41, 42, and 41, respectively (Table S1). Average age of Ashkenazi-Jewish-European and African ancestry donors was higher (average 53 and 54, respectively). None of these differences can be attributed to differences in proliferation rate of cells in the breast during tissue collection as MKI67, a marker of cell proliferation, is expressed in very few cells across samples (Figure 2e).

**Figure 2:**
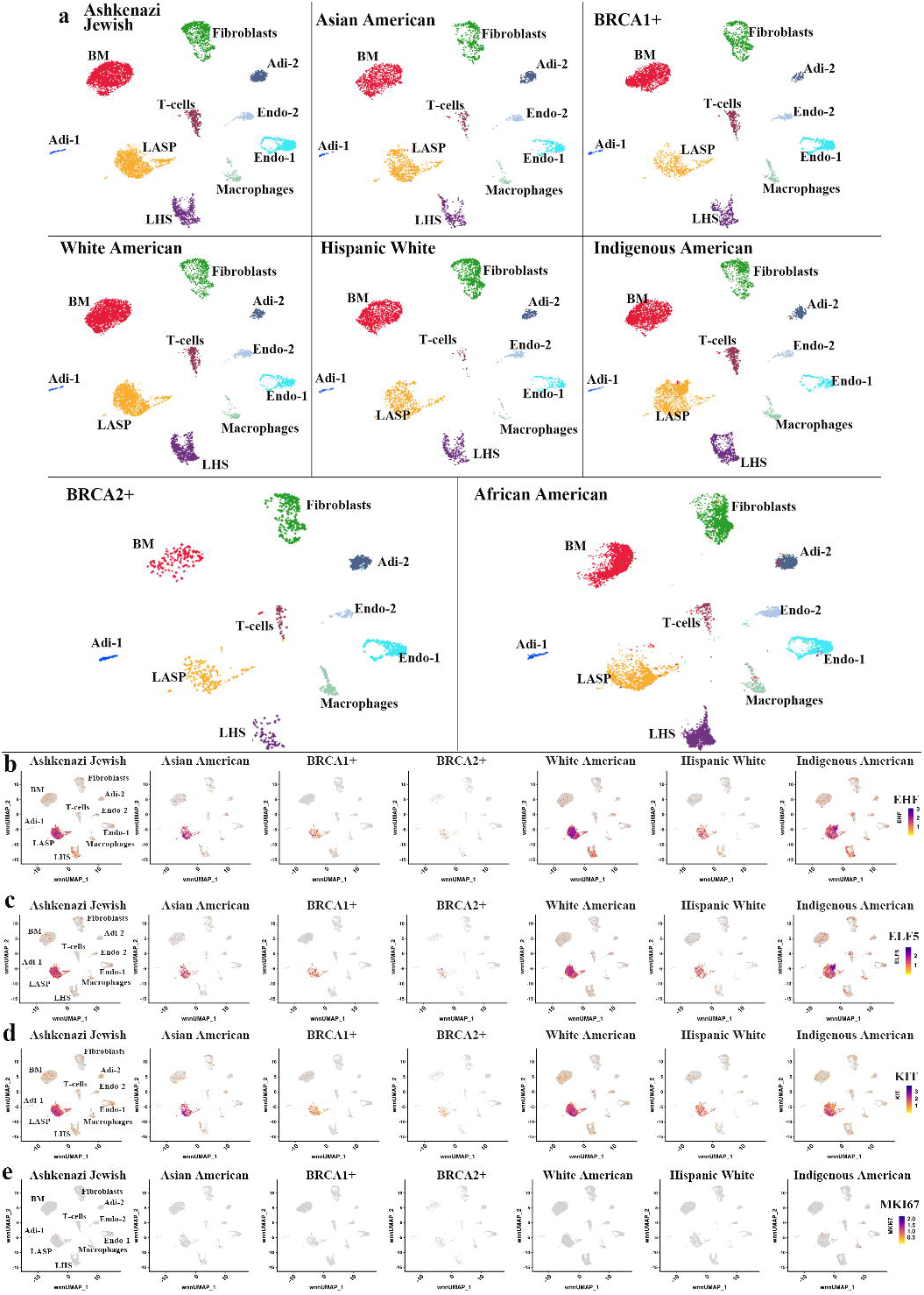
Genetic ancestry-dependent variability in cell state. a) Cell clustering in each group based on integrated snATAC-seq and snRNA-seq analyses. b) Expression pattern of alveolar cells marker EHF. c) Expression pattern of alveolar cells marker ELF5. d) Expression pattern of luminal progenitor cells marker KIT. e) Expression pattern of the cell proliferation marker MKI67.

### AP cells enriched in Indigenous Americans express ESR1 and are enriched for transcripts downstream of ER-growth factor signaling

To determine whether AP cells enriched in Indigenous Americans express a unique set of genes compared to other cell clusters within LASP cells, we compared gene expression between cluster 9 (Indigenous American enriched cluster) and clusters 1, 12 and 18 (Table S5). ESR1 is among the top genes highly expressed (∼230 fold higher) in this cluster compared to other clusters. ESR1 expression pattern in various clusters, as shown in Figure 3a, further confirmed these results. A set of pioneer factors control ERα genome wide binding including FOXA1 and GATA3 and ERα-FOXA1-GATA3 constitute a cell type specific transcription factor network of hormone responsive cells ^17,18^. FOXA1 expression was mostly restricted to LHS cells, even in Indigenous Americans, whereas GATA3 expression was higher in LHS cells with lower level expression in LASP and BM cells (Figure 3b and Figure 3c).

**Figure 3:**
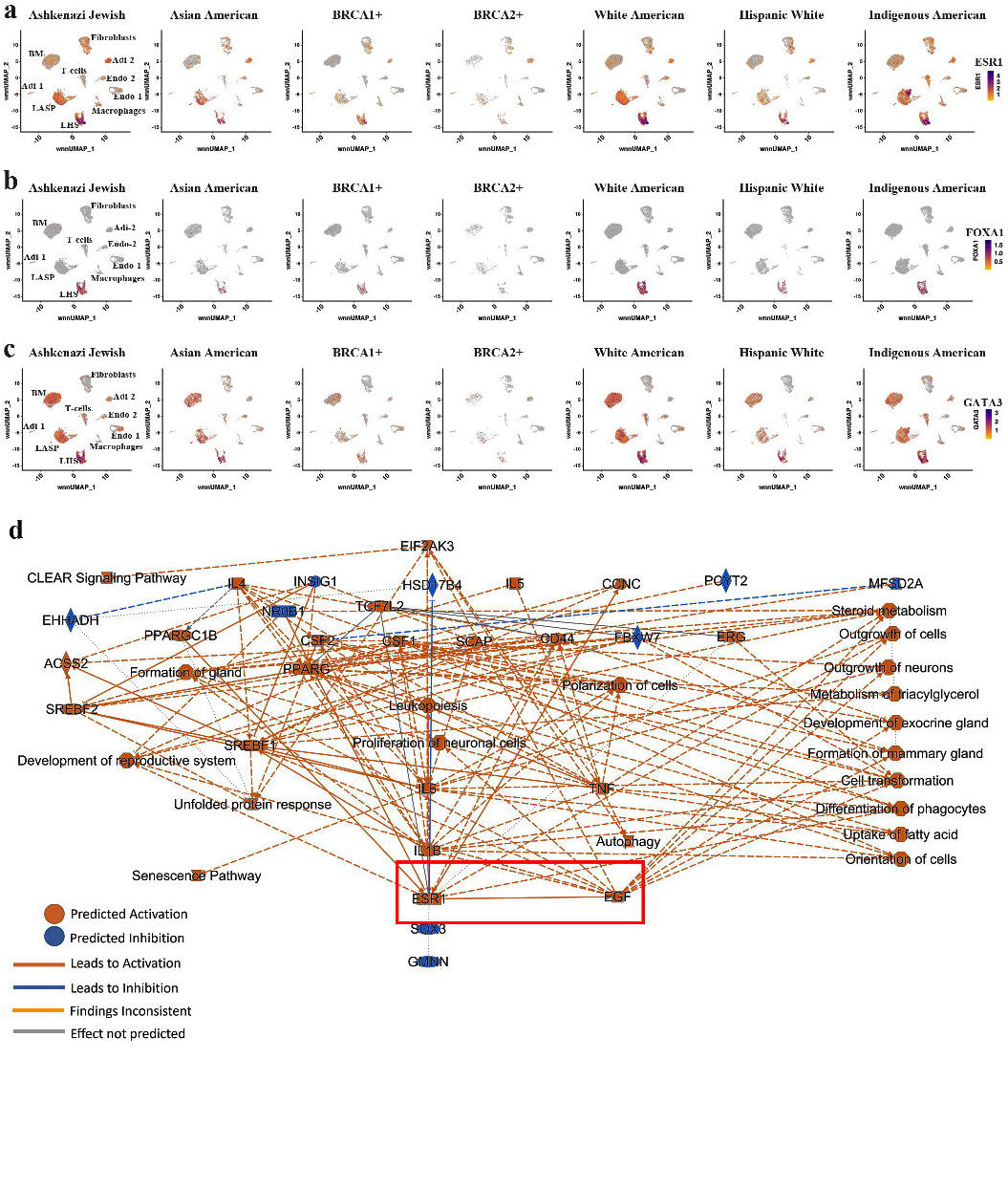
Genetic ancestry-dependent variability of ESR1, FOXA1, and GATA3, which constitute a hormone responsive cell lineage enriched transcription factor network. a) Expression pattern of ESR1. b) Expression pattern of FOXA1. c) Expression pattern of GATA3. d) AP cells enriched in Indigenous Americans show elevated ER-growth factor signaling crosstalk. Genes differentially expressed in AP cells compared to other cell state among LASP cells were subjected to IPA and the top signaling network is presented.

Lack of FOXA1 expression but the presence of ESR1 transcripts in AP cells of Indigenous Americans suggested that ERα activity in these cells is regulated by a mechanism distinct from that in LHS cells. To evaluate this possibility, we subjected genes differentially expressed in cluster 9 to Ingenuity Pathway Analysis. Among the various pathways enriched in this cluster, intersection of EGF signaling with ER signaling emerged as one of the top signaling networks (Figure 3d). EGF has previously been shown to alter the ER cistrome and induce gene expression patterns found in antiestrogen resistant cells ^19^. Thus, in cluster 9, ER and EGF signaling cross talk may be dominant. To further confirm this possibility, we next determined how many of the genes differentially expressed in cluster 9 are known ER-regulated genes in the breast cancer cell line MCF-7 ^20^. We determined overlap between cluster 9 enriched genes and our previously described ERα:E2 regulated genes in MCF-7 cells ^20^. Among 671 genes enriched in Cluster 9, 264 genes are ERα:E2 regulated genes (Table S13). Therefore, it is possible that E2 and growth factors control ERα activity in LHS and LASP cells, respectively.

### Chromatin accessibility of ESR1 gene extends to LASP cells

To determine whether cell type specific expression of ESR1 correlates with chromatin accessibility, we mapped chromatin accessible regions of ESR1 in various epithelial cell types and among donors of different genetic ancestry. Accessibility map included 10 KB regions upstream of transcription start site to cover promoter regions and 10 KB downstream of transcription end site to cover potential 3’ enhancer regions. Although ESR1 expression was limited to LHS cells among most donors except Indigenous Americans, the ESR1 gene displayed similar chromatin accessible regions in both LHS and LASP cells (Figure 4a) except for one peak (peak 1) being more prominent in LHS cells. Since ESR1 is transcribed from multiple promoters ^21^, it is difficult to assign which among the open chromatin peaks correspond to promoter and enhancers. Since coding region of one of the ESR1 isoforms (NM_000125) starts at chr. 6:151,807,682, peaks 2 and 3 can be considered open chromatin regions of enhancer-promoters. Peak 4 may correspond to promoter region for isoforms such as ERαΔ36, which is transcribed from an intronic promoter ^22^. There was genetic ancestry and BRCA2 mutation dependent variability in chromatin accessible peaks 1 and 4 (Figure 4b). For example, peak 4 is more prominent in BRCA2 mutation carrier but close to being absent in Indigenous Americans. ESR1 binding sites are present in chromatin accessible regions of LHS and LASP cells but at lower levels in BM cells, although differences are less dramatic (Figure 4c). With respect to FOXA1, promoter region was more accessible in LHS cells compared to other cell types (Figure 4d). Furthermore, chromatin accessible regions of LHS cells were enriched for binding sites for FOXA1 compared to other cell types (Figure 4e).

**Figure 4:**
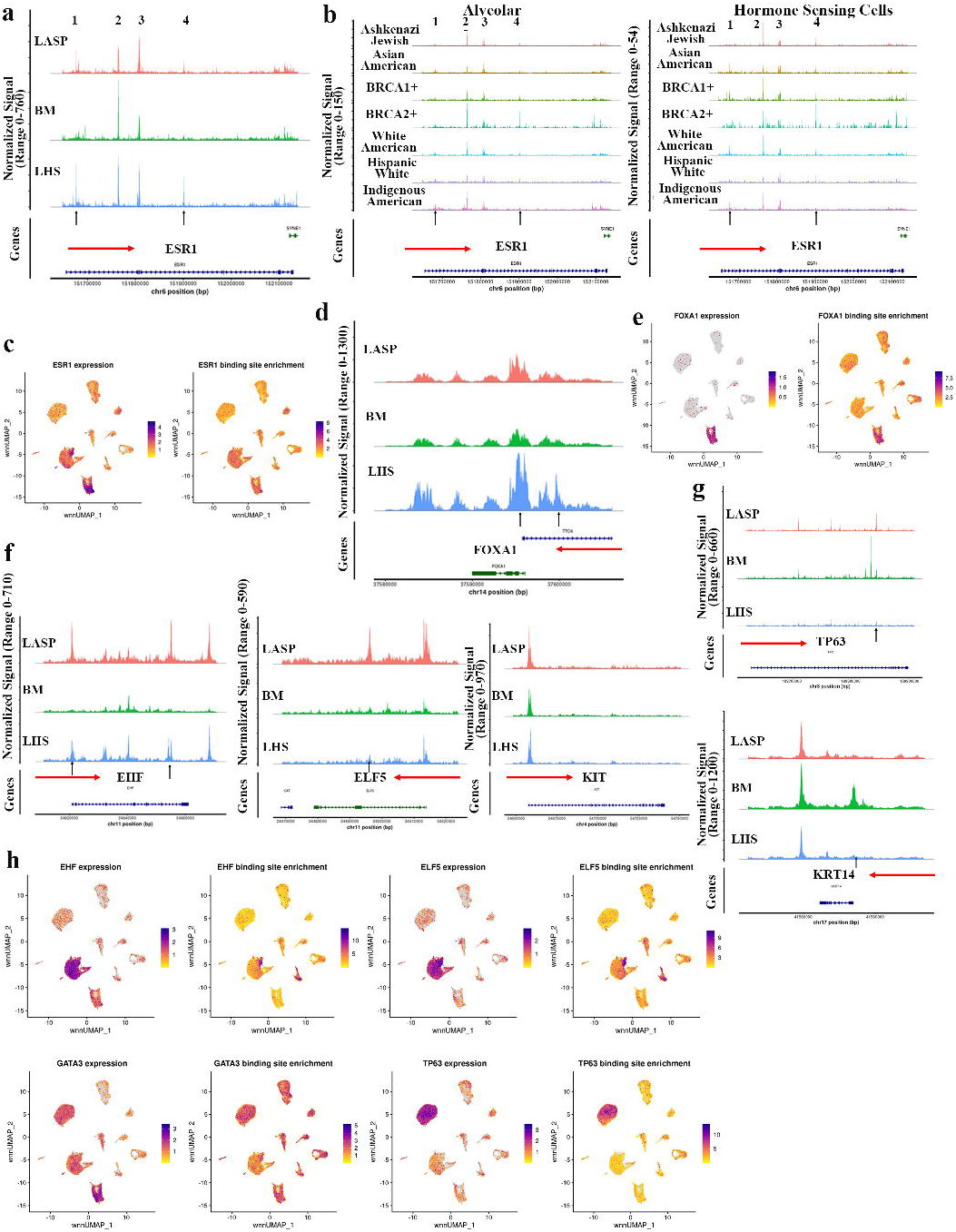
Relationship between chromatin accessibility and gene expression. a) ESR1 chromatin accessibility patterns in LSH, LASP, and BM cells. Horizontal red arrow marks the direction of the indicated gene transcription. Vertical arrow denotes cell type-specific chromatin accessible regions. b) Chromatin accessibility map of ESR1 gene in LHS and LASP cells of breast tissues of women of different genetic ancestry. The chromatin accessible peaks are numbered 1-4 and few of these peaks showed genetic ancestry and BRCA2 mutation status dependent variabilities. c) Binding sites for ER are present in the chromatin accessible regions of multiple cell types, including cell types in which ESR1 is not expressed. d) Chromatin accessibility map of FOXA1 in various epithelial cell types of the breast. e) Binding sites for FOXA1 in chromatin accessible regions of various cell types. f) Chromatin accessibility map of LASP markers EHF, ELF5, and KIT in various cell types of the breast. g) Chromatin accessibility map of BM cell markers TP63 and KRT14 in various epithelial cell types of the breast. h) EHF, ELF5, GATA3, and TP63 expression patterns and binding site enrichment analysis.

We also examined the chromatin accessibility status of LASP/alveolar markers EHF, ELF5, and KIT. Distinct chromatin accessibility patterns for EHF and ELF5 were observed in three major epithelial cell types of the breast with prominent chromatin accessible promoter region in LASP cells followed by LHS cells (Figure 4f). However, promoter regions of KIT was similarly accessible in all three cell types. BM cells had least open chromatin regions for these two genes. Therefore, EHF and ELF5 are more reliable markers of LASP cells. Chromatin accessibility map of BM cell markers TP63 and KRT14 showed cell type-dependent variability with BM cells showing distinct chromatin accessible promoter region (Figure 4g). Multiple isoforms of TP63 are expressed using different promoters ^23^ and it is difficult to ascertain which among the chromatin accessible regions contain cell-type specific active promoter/enhancer regions of TP63. Nonetheless, these results clearly show that TP63 and KRT14 are bona fide markers of BM cells and changes in their expression in disease conditions require chromatin reorganization.

Expression patterns and binding site enrichment analysis showed that binding sites for EHF and ELF5 are enriched in the AP subpopulation of LASP cells (Figure 4h). TP63 expression and binding site enrichment were restricted to BM cells, whereas GATA3 expression and binding site enrichment did not show cell type specificity (Figure 4h). Collectively, only select genes expressed in epithelial cell types of the breast show cell type-specific chromatin accessibility patterns and their cell type-specific expression is influenced likely by extracellular signals rather than at the level of cell type specific chromatin accessibility.

### Incompatibility in gene expression and chromatin accessibility extends to other genes associated with cell identity

Several in vitro studies have shown that the transcription regulator ZEB1 mediates epithelial to mesenchymal transition of breast epithelial and breast cancer cells and that its promoter region remains in poised state to confer plasticity to breast cancer cells ^24–26^. However, our previous two studies examining ZEB1 protein levels in normal breast tissues demonstrated the presence of ZEB1^+^ cells in stroma that surrounds duct with no expression in epithelial cells ^13,27^. We examined our multiome dataset for ZEB1 expression and chromatin accessibility. Interestingly, ZEB1 mRNA expression was observed in fibroblast-like cells, endothelial cells, adipocytes, and T cells but not in epithelial cells (Figure 5a). However, ZEB1 regulatory regions showed similar patterns of chromatin accessibility in fibroblasts and all three epithelial cell types (Figure 5b).

**Figure 5:**
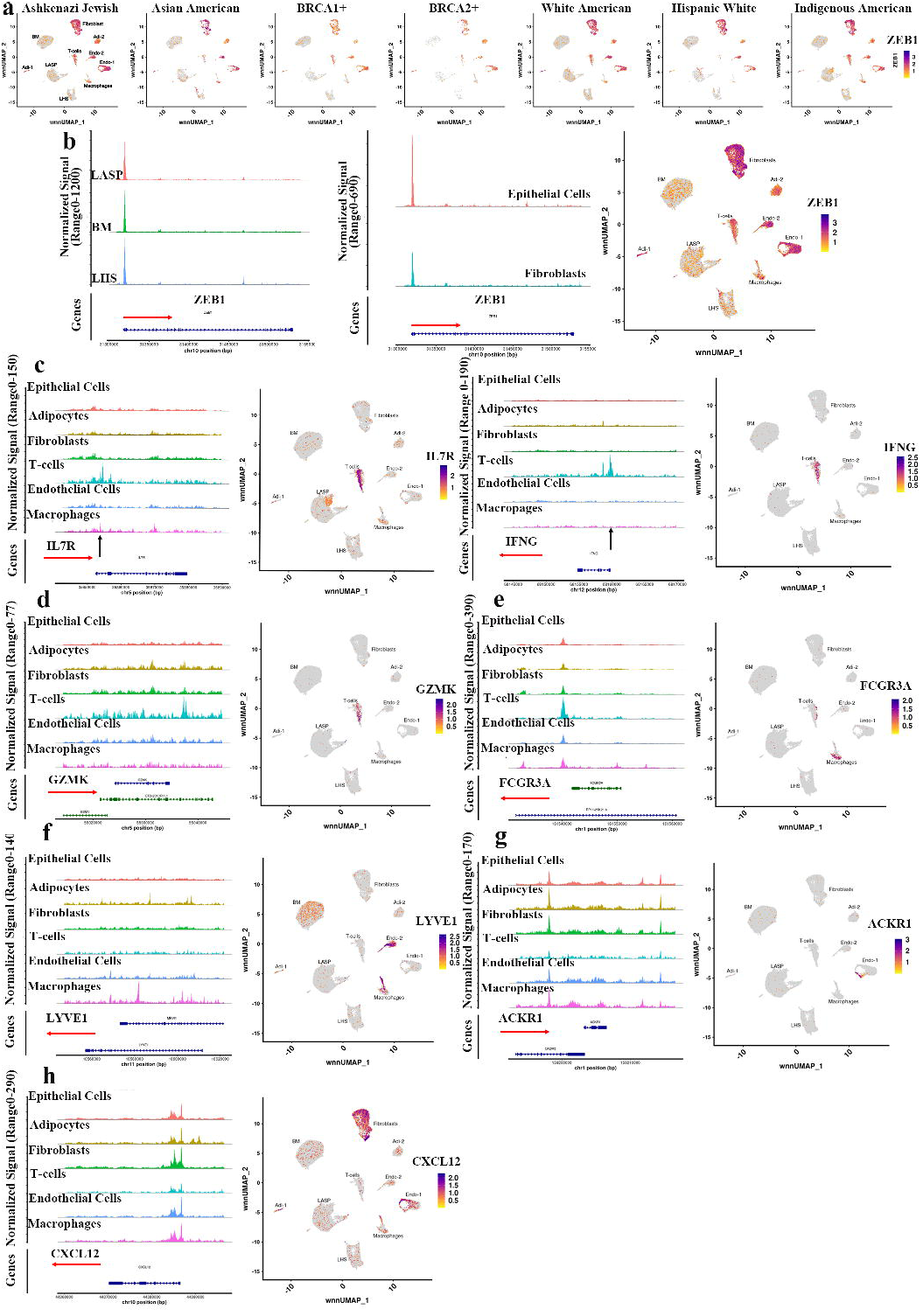
Limited relationship between cell-type specific gene expression and chromatin accessibility. a) Epithelial cells do not express ZEB1. b) Despite fibroblast-restricted expression, chromatin accessibility of ZEB1 is similar between fibroblasts and epithelial cells. c) IL7R and IFNγ expression and chromatin accessibility are restricted to T cells. d) GZMK expression and chromatin accessibility are restricted to T cells. e) FCGR3A expression and chromatin accessibility are restricted to macrophages. f) Lymphatic endothelial marker LYVE1 is expressed in Endo-2 and a fraction of macrophages but the chromatin accessibility patterns were not unique to these two cell types. g) Although ACKR1 expression is restricted to a subpopulation of endothelial cells, ACKR1 gene showed limited variation in chromatin accessibility between various cell types. h) CXCL12 expression and chromatin accessibility showed limited correlation.

We extended the above observations to other genes that are used as markers to define various endothelial and T cell subtypes. For example, CD4^+^ T cell enriched IL7R and CD8^+^ T cell enriched IFNγ showed T cell-specific expression and chromatin accessibility (Figure 5c). Similarly, expression and chromatin accessibility of CD8+ T cell enriched GZMK were restricted to T cells (Figure 5d). Chromatin accessibility of macrophage enriched FCGR3A was restricted to macrophages (Figure 5e).

Among the endothelial cells, lymphatic endothelial cell marker LYVE1 is expressed in Endo-2 subcluster and a macrophage subpopulation but the chromatin accessibility patterns in these two cell types were not similar (Figure 5f). Similarly, Endothelial Stalk-like subtype marker ACKR1 is expressed in a subset of Endo-1 subcluster but the chromatin in the regulatory regions of this gene is accessible in all cell types (Figure 5g). CXCL12, which is expressed in Endo-1 and in fibroblasts, showed similar chromatin accessible patterns in all cell types except T cells (Figure 5h). These results collectively demonstrate lack of compatibility in chromatin accessibility and expression of corresponding genes in the majority of cases.

### Breast tissues of women of African ancestry show fibroblast and epithelial cell states distinct from breast tissues of women of European ancestry

As noted above, our multiple attempts to perform integrated snRNA-seq and snATAC-seq of breast tissues of women of African ancestry were not successful. Since breast cancer outcome disparity is known among African American women and our prior studies have suggested a biologic basis of disparity, particularly related to composition of stromal cells ^3,13,27^, we made multiple attempts through different protocols but were finally successful in obtaining reliable data only when nuclei were subjected to snRNA-seq alone. To allow appropriate comparison, we subjected new breast tissues of women of European ancestry to a similar protocol and performed snRNA-seq. Cell clustering showed similarity to clustering patterns obtained with integrated snATAC-seq and snRNA-seq (Figure 6a). However, there were major differences in epithelial and fibroblast cell states between breast tissues of women of African and European ancestry. For example, LASP cell clusters in African ancestry are dominantly populated by BL cell state, whereas in case of European ancestry, there were similar numbers of BL and AP cells. Similar to data presented in Figure 3a, ESR1 and FOXA1 expression were restricted to LSH cells in both groups (Figure 6b). Unlike in the breast tissues of Indigenous Americans, AP cells did not express higher levels of ESR1 compared to other LASP cells. Average age of donors of African ancestry and European ancestry in this snRNA-seq was 54 and 34, respectively. Therefore, we compared cell type status of women of African ancestry with that of the Ashkenazi-Jewish European group with similar age (average 53). Differences between African and European ancestry persisted when cell state of African Ancestry was compared to cell state of Ashkenazi-Jewish European group as shown in Figure 2a. The differences between cell clusters of women of African and European ancestry are not due to differences in proliferation rate as there was only a minor difference in MKI67+ cells between two groups (Figure 6c) and MKI67 positivity was similar between African and Ashkenazi Jewish-European groups.

**Figure 6:**
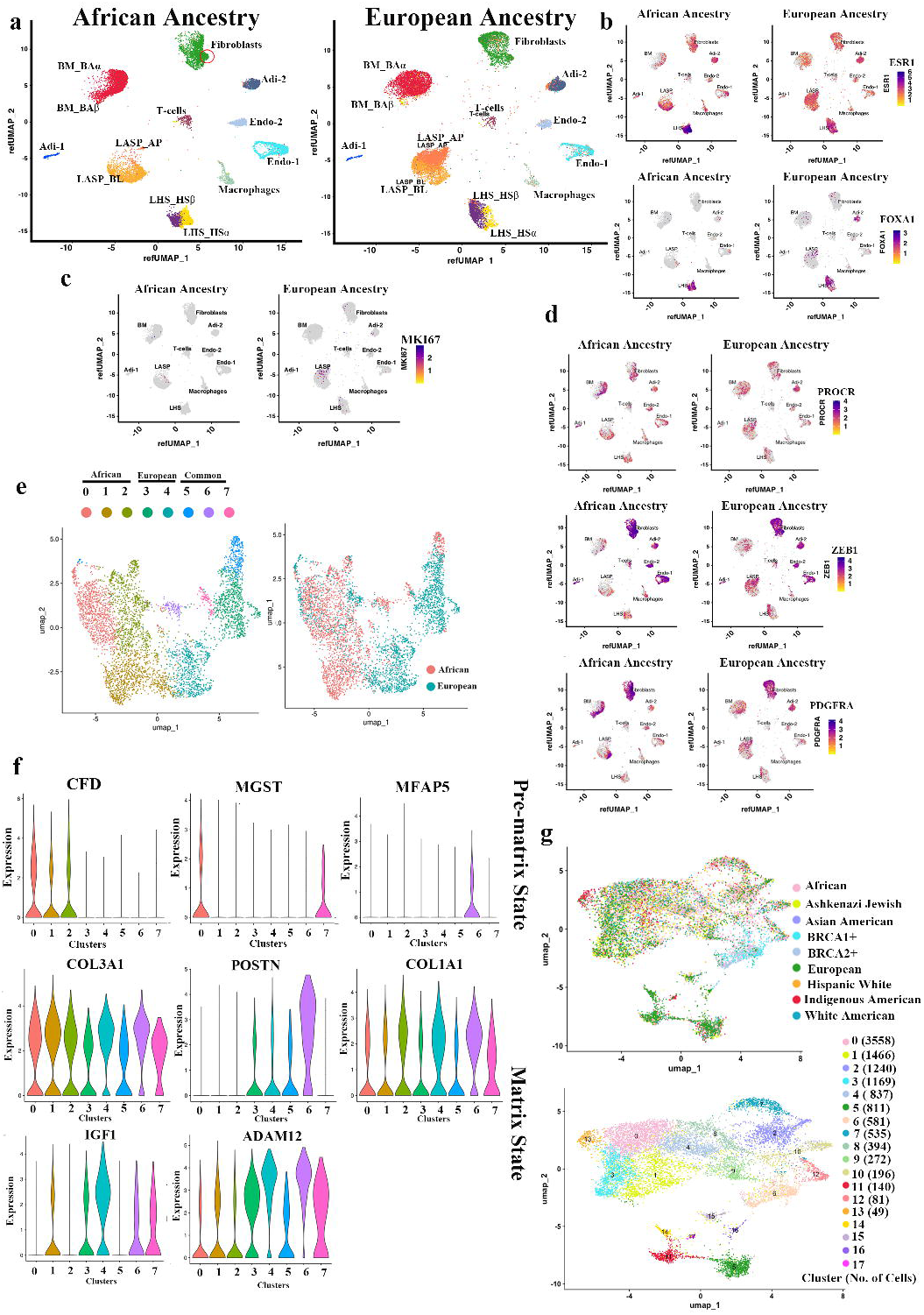
Comparative analyses of breast tissues of women of African ancestry with European ancestry using snRNA-seq. a) Fibroblasts and epithelial cells of the breast tissue cluster differently in African ancestry compared to European ancestry. b) ESR1 and FOXA1 expression patterns in epithelial cell clusters of African and European ancestry. c) MKI67 expression patterns in breast tissues of African and European ancestry. d) PROCR, ZEB1 and PDGFRα expression patterns in breast tissues of African and European ancestry. d) Fibroblasts in African and European ancestry show distinct cell states. e and f) Fibro-prematrix state dominate in African ancestry, whereas Fibro-matrix state dominate in European ancestry. g) Genetic ancestry- and germline mutation-dependent variability in clustering of fibroblasts.

We have recently reported a unique type of stromal cells with multi-potent activity enriched in breast tissues of women of African ancestry ^13^. These cells are PROCR^+^/ZEB1^+^/PDGFRα^+^ (hence called PZP cells) and display mesenchymal stem and fibroblast-like features. Breast epithelial cells co-cultured with PZP cells transdifferentiated into BL state ^13^, which may explain the elevation of BL-like LASP cells in the breast tissues of women of African ancestry. Because fibroblasts in breast tissues of women of African ancestry clustered differently from that of European ancestry, we analyzed the expression patterns of PROCR, ZEB1 and PDGFRα. A subpopulation of cells in the fibroblast clusters were positive for all three markers and, number of PROCR+/ZEB1+/PDGFRα+ cells were higher in fibroblast cluster of women of African ancestry compared to women of European ancestry (Figure 6d, indicated by a circle in Figure 6a), consistent with our prior study showing protein level differences ^13^.

Fibroblasts in breast have recently been shown to exist in four different states based on expression pattern of specific genes-Fibro-prematrix, Fibro-SFRP4, Fibro-major and Fibro-matrix ^7^. Using the gene expression signatures that specify these cell states, we analyzed fibroblasts of breast tissues from different genetic ancestry donors. As shown in Figure 6e, there were genetic ancestry-dependent variabilities in fibroblast state with women of African ancestry showing specific enrichment of genes that specify Fibro-prematrix state at the expense of Fibro-matrix state (Figure 6e and 6f). Figure S2 provides additional differences in fibroblasts between African and European ancestry and gene expression differences between fibroblasts of African ancestry and European ancestry are shown in Table S14. It is interesting that the expression of several of the ATP binding cassette subfamily members is elevated in fibroblasts of African ancestry compared to European ancestry (Figure S2). We also note that none of the markers corresponding to other fibroblast clusters are expressed at high levels. (Figure S2). When fibroblast populations of other ancestry groups and BRCA1/2 mutation carriers were compared, there was considerable variation in fibroblast cell state across genetic ancestry and BRCA1/2 mutation status suggesting important contribution of fibroblast diversity in human health (Figure 6g). For example, consistent with the recent report on the effects of BRCA1 mutation on stromal cells ^28^, fibroblasts in BRCA1/2 mutation carriers are enriched in cluster 6.

Previous studies have demonstrated that breast tumors in African Americans are enriched for exhausted T cells ^29^. To determine whether healthy breast tissues of women of African ancestry intrinsically contain higher levels of exhausted T cells, we determined the expression patterns of CD274 (PD-L1), CTLA-4, LAG3, and PDCD1, all markers of exhausted T cells, in breast tissues from women of African and European ancestry ^29^. No significant differences were noted (Figure S3) between tissues of African and European ancestry. Therefore, accumulation of exhausted T cells in tumors of African American women is likely induced by the tumor microenvironment.

Murrow et al recently reported hormone-regulated cell-cell interaction network in the human breast at single cell resolution and identified various markers that can distinguish hormone responsive cells into two states; LRRC26 marks cells with both ER and progesterone receptor activity (HR+ State 1), whereas P4HA1^+^ cells to represent hormone responsive cells with elevated hypoxia/Pro-angiogenic activity (HR+ State 2) ^9^. Although LRRC26 expression was seen in a subpopulation of LHS cells, a subset of LASP cells also expressed this gene (Figure S4). Every cell type of the breast expressed P4HA1, suggesting lack of specificity of this gene as a marker of hormone responsive cells. We extended this analysis to include other genes suggested to be enriched in HR+ State 1 (PNMT, CXCL13, MYBPC1, CADP52, WNT4, PPP1R1B, and IL20RA) and HR+ State 2 (NDRG1, HILPDA, ANGPTL4, EGLN3, ERO1A, PLOD2, and ENO2). Only a few of these genes (CXCL13, MYBPC1, PNMT and WNT4) are expressed at higher levels in ESR1-positive cells (Figure S4). Additional studies are needed to further characterize distinct hormone responsive cell states.

### Individual level characterization of age-dependent changes in breast epithelial cell gene expression

Several donors to our tissue bank have donated breast tissues twice, 10 years apart (described as timepoint 1 and timepoint 2) allowing us to determine gene expression differences with age. Prior studies focused on identifying age-dependent differences in gene expression utilized tissues from independent donors of different ages ^8^. Because of limited tissue samples, we performed spatial transcriptomics with samples from five donors who donated breast tissues twice. However, consistent results covering all regions of interest (ROI) were obtained with three pairs of samples. Age and BMI status of three donors at both times of tissue collection, UMAP of transcriptome data, and immunostaining of breast tissues to differentiate epithelial cells and adipocytes are shown in Figure S5a and b). Micro-dissected ducts and lobules as well as adipocytes are shown in Figure 7a. We first evaluated resulting spatial transcriptomics data for adipocyte, endothelial and epithelial cell subtype gene signatures derived from multiome data (Tables S4-S9). These initial analyses showed that data from timepoint 2 cluster into two groups, but are generally more abundant in adipocyte-2 (Adi-2), macrophages and Endo-2 cell types compared to timepoint 1 (Figure S5c). Adi-1 and Adi-2 differ with respect to adiponectin expression with Adi-1 expressing 30-fold higher adiponectin than Adi-2 (Table S9). Endo-2 is enriched for lymphatic endothelial cell markers. Spatial transcriptomics allowed us to extend the studies to find gene expression differences between lobular and ductal epithelial cells. Volcano plot in Figure 7b shows differences in gene expression between ductal and lobular epithelial cells. Pathway analysis of genes differentially expressed in ductal and lobular epithelial cells and at two different time points are shown in Figure S6. While metabolic pathways are enriched in epithelial cells of the lobules, ductal epithelial cells showed enrichment of extracellular signal activated signaling networks such as cytokine and cancer-associated signaling networks. Age-dependent changes were also restricted to metabolic pathways in lobular epithelial cells, whereas changes in splicing machinery were observed in ductal epithelial cells. Genes differentially expressed in lobular epithelial cells compared to ductal epithelial cells (Table S15), differences at timepoint 1 (Table S16), at timepoint 2 (Table S17); within ducts between timepoint 1 versus 2 (Table S18), within lobules between timepoint 1 versus 2 (Table S19) are provided as supplementary information. Volcano plots shown in Figure 7c-7f further highlight key genes differentially expressed in ductal and lobular epithelial cells under different conditions. Four-hundred twenty six genes are differentially expressed in ductal versus lobular epithelial cells. A recent study using a different spatial transcriptomic technique listed 30 genes that are expressed differentially in ductal and lobular epithelial cells ^7^. Among those 30 genes, 10 genes showed similar expression patterns (MGP, ANXA1, TACSTD2, KRT14, KRT17, WFDC2, STAC2 and ALDH1A3 are elevated in ductal epithelial cells whereas APOD and SNORC are elevated in lobular epithelial cells). Expression pattern of these genes assessed through multiome data is shown in Figure S7. Since 10 genes showed similar expression patterns in lobular and ductal epithelial cells in two independent studies, these genes can be considered as markers of ductal and lobular epithelial subtypes. However, expression of only few of these genes is enriched in epithelial cells. We observed upregulation of KRT14 and KRT17, BM cell enriched genes, in ductal compared to lobular epithelial cells, consistent with the previous report ^7^. Immunoglobulin heavy constant alpha 1 (IGHA1) and Immunoglobulin Kappa Constant (IGKC) gene, both of which are expressed in the breast as per GTEx database, are expressed at higher levels in lobular compared to ductal epithelial cells. Several genes showed differences in expression in two different timepoints but only PTBP1 showed consistent decline (∼six folds) in expression in timepoint 2 compared to timepoint 1 in both ductal and lobular epithelial cells. PTBP1 is associated with mRNA processing and alternative splicing ^30^ and, based on UALCAN database ^31^, PTPB1 is overexpressed in all subtypes of breast cancer (Figure S8). We determined genes commonly upregulated or down-regulated at timepoint-2 compared to timepoint-1 in both lobular and ductal epithelial cells. IPA analyses of the genes showed downregulation of genes associated with PKA signaling pathway but upregulation of EIF2 and oxidative phosphorylation pathways in timepoint 2 compared to timepoint 1 (Figure 8 and Table S20).

**Figure 7:**
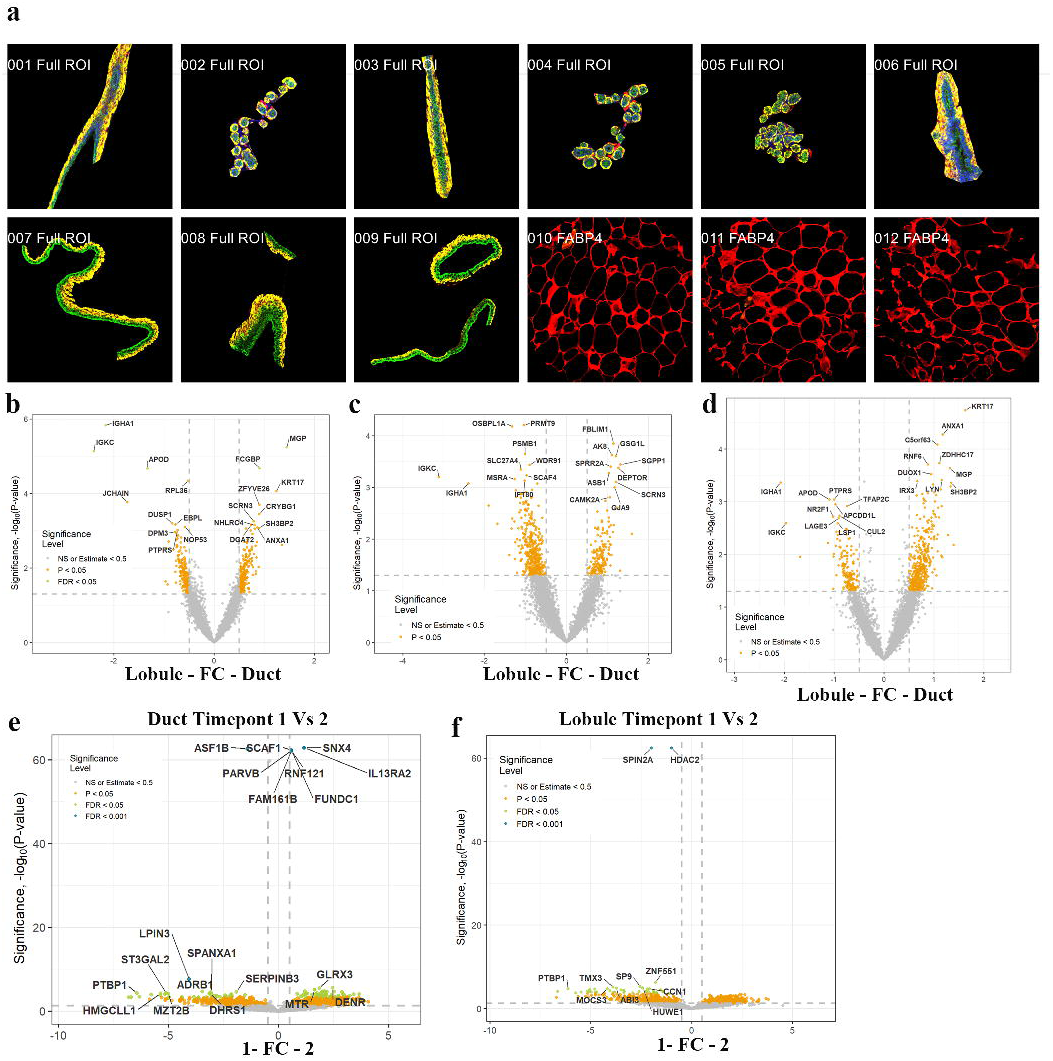
Spatial transcriptomics reveal gene expression differences between ductal and lobular epithelial cells. To avoid inter-individual variation, few of the plots are from donor #3. a) Images of micro-dissected ducts, lobules and adipocytes. b) Volcano plot showing gene expression differences between lobular and ductal epithelial cells. c) Gene expression differences between ductal and lobular epithelial cells at timepoint 1. d) Gene expression differences in ductal and lobular epithelial cells at timepoint 2. e) Gene expression differences in ductal epithelial cells between timepoints 1 and 2. f) Gene expression differences in lobular epithelial cells between timepoints 1 and 2.

**Figure 8:**
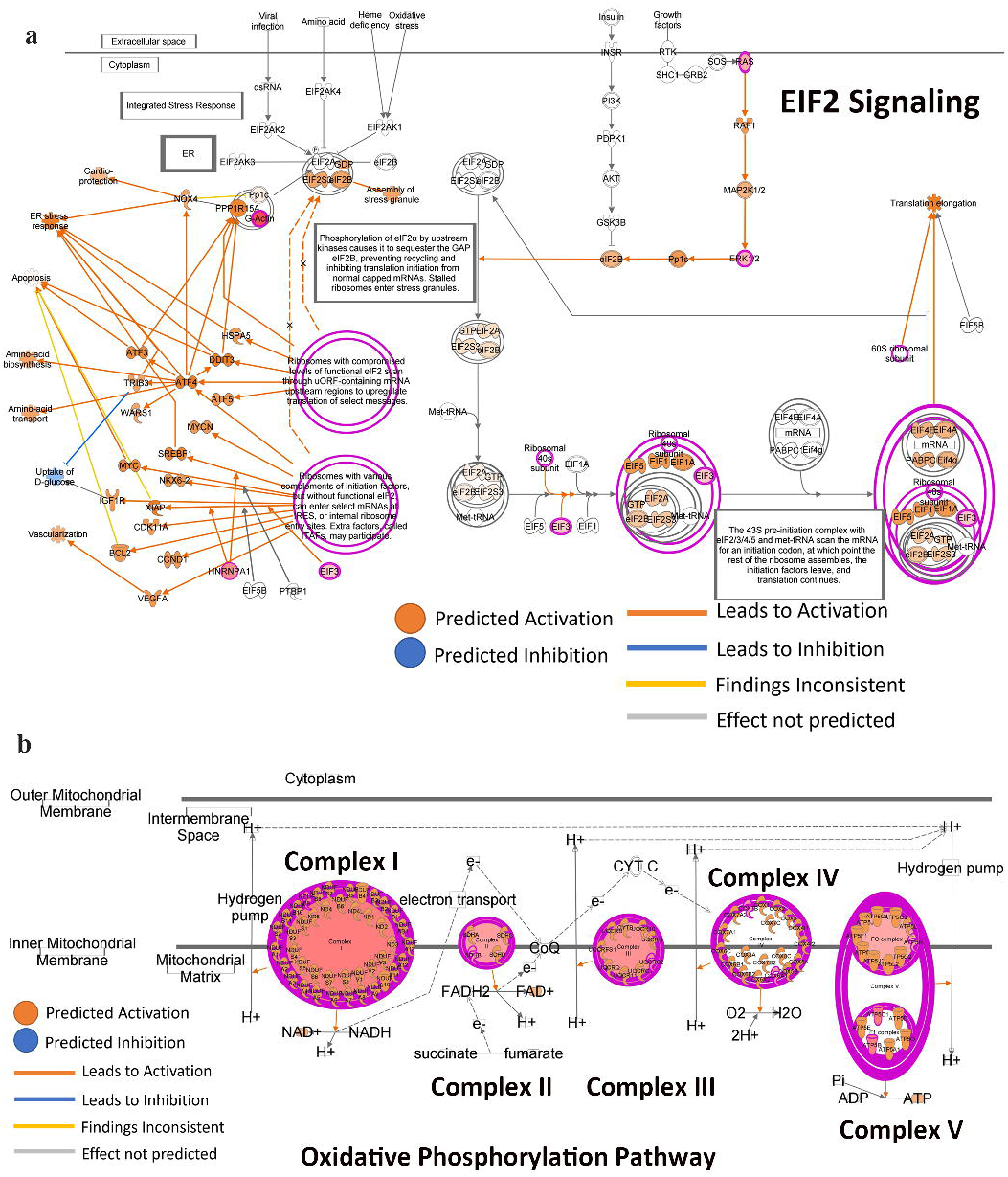
EIF2 and oxidative phosphorylation pathways are upregulated but PKA pathway is downregulated at timepoint 2 compared to timepoint 1 in breast epithelial cells.

## Discussion

The biomedical research community has recognized the importance of including tissues from people of different genetic ancestry to make research representative of all people ^32^. This recognition came from several clinical observations highlighting genetic ancestry-dependent variability in utility of commonly used disease biomarkers, mutation patterns in cancer, response to treatment, and ultimately disease outcome ^6,33,34^. For example, a recent study showed that genetic ancestry but not the amount of UV exposure dictates clonal architecture and skin cancer susceptibility ^35^. To aid disease specific research, global efforts such as Human Cell Atlas and Chan-Zuckerberg initiatives are focused on developing a single cell atlas of the body. It is recognized by these organizations that reference a single cell atlas of the body should also include tissues from people of diverse genetic ancestry. This study was undertaken to meet this goal for the breast tissue atlas. By sequencing 88,005 nuclei from breast tissues collected using the same standard operating procedure (details are in Komen Tissue Bank website) from women of diverse genetic ancestry and with no clinical history of breast cancer, we have developed a global breast single cell atlas. Unlike other efforts in this direction, which used reduction mammoplasty or normal tissues adjacent to tumors, which we and other have shown to be histologically abnormal ^27,36,37^, our study utilized tissues from healthy donors. Because we subdivided the samples for the analyses based on self-reported race/ethnicity and verified further with genetic ancestry analysis, we were able to present a single cell atlas of different groups based on genetic ancestry and to identify two major differences based on ancestry-alveolar progenitor cells in Indigenous Americans and stromal fibroblasts in women of African ancestry. There have been many attempts to understand the biologic basis of breast cancer disparity based on genetic ancestry. Most of these studies focused on identifying differences in genomic aberrations in cancer cells and immune cells that infiltrate the tumors ^6,29,38^. However, genetic ancestry-dependent differences in stromal cells such as fibroblasts could be another source of disparity as we demonstrated recently ^13^ and need further attention.

A distinct difference in ESR1 expression in breast epithelial cells of Indigenous Americans compared to others is interesting and alveolar progenitor cells that express ESR1 in Indigenous Americans have a gene expression program that is influenced by crosstalk between estrogen receptor and growth factor signaling. Recent studies have suggested that Indigenous Americans are more susceptible to ERBB2+ breast cancers ^4^. It is unknown whether distinct gene expression program in alveolar cells that involves ER-growth factor crosstalk increases susceptibility to ERBB2+ breast cancers. ESR1 upregulation in alveolar progenitor cells of Indigenous Americans is not due to distinct chromatin organization but rather appears to be at the level of transcriptional output. In this respect, a breast cancer protective allele found mostly in Indigenous Americans has been suggested to affect ESR1 expression ^5^. However, that protective allele is less likely to be responsible for our observation as only two out 10 donors carried this protective allele. Several other variants of ESR1 linked to various conditions have been described in the literature ^39–41^ and it is possible that a few of these variants affecting ESR1 expression or transcript stability are more prevalent in Indigenous Americans.

Our studies revealed that FOXA1 but not ESR1 is a reliable marker of LHS cells as its expression was much more restricted to LHS cells than any other genes we examined. Furthermore, FOXA1 but not ESR1 showed unique chromatin accessibility patterns in LHS cells. In our previous studies, we had observed FOXA1 expression in 300 out of 404 breast tumors and three cell states among LHS cells are putative cells-of-origin of breast cancer ^11,42^. Thus, it is likely that the majority of breast cancers, particularly ER+ breast cancers, originate from LHS cells instead of LASP cells.

Breast cancer outcomes in women of African ancestry is generally poor even after adjusting for socioeconomic factors and healthcare access ^3^. The biologic basis for this disparity is just beginning to be elucidated. Genetic ancestry-dependent variability in the tumor microenvironment, particularly immune cells, and inflammatory cytokines has been reported ^29,43^. We reported enrichment of stromal cells with mesenchymal stem-like and fibroadipogenic properties in the breast tissues of women of African ancestry and the ability of these cells to alter cytokine/chemokine profiles ^13^. Fibroblast subcluster described in this study also show distinct cell state in women of African Ancestry compared to others. Fibroblast cell state in women of African ancestry is predominantly prematrix state compared to others and this cell state is associated with vasculogenesis ^7^. How this fibroblast state impacts the normal biology as well as breast tumor biology remains to be investigated. Considering genetic ancestry-dependent diversity fibroblast state observed in this study, understanding fibroblast biology will likely provide new insight into biologic basis of health disparity.

Integrated snATAC-seq and snRNA-seq revealed that chromatin accessibility is not the primary determinant of cell type-specific expression of most of the genes in the breast. As breast tissue undergoes frequent remodeling in response to hormonal cues during puberty, menstrual cycle, pregnancy, lactation, involution, and menopause, ability of cells to respond to extracellular signals rapidly and plasticity are very much essential during remodeling. Gene expression changes that are not dependent on changes in chromatin accessibility are required in these situations. In this context, it is interesting that while ESR1 expression is restricted to LHS cells of the breast of women of all but Indigenous American ancestry, chromatin accessibility allowing gene expression is maintained in both LHS and LASP cells with minor differences. Genomic aberration in any transcription factor that controls ESR1 expression could lead to ESR1 expression in LASP cells and make these cells responsive to estrogen and potentially promote carcinogenesis. Several transcription regulators such as GATA3, FOXA1, EP300, TCF7L2, TFAP2C, BRCA1, TP53 and BARX2 control the expression of ESR1 ^21^ and cBioportal analyses revealed in at least 50% of breast cancers one or more of these transcription regulators show genomic aberrations. ESR1 gene appears to be organized differently in BRCA2 mutation carriers than others. BRCA2 mutation carriers typically develop ER+ breast cancers ^44^ and whether distinct organization of ESR1 gene in these carriers leads to altered ESR1 isoform expression remains to be determined. Once a human single cell atlas of multiple organs is developed, it will be interesting to determine cells in how many organs have cells that maintain such a relationship between gene expression and chromatin accessibility, and whether cancer susceptibility of an organ is linked to a relationship between gene expression and chromatin accessibility.

Breast cancers originating from epithelial cells of the duct and lobules show distinct biology, clinical features, and response to treatment ^45^. Therefore, there has been considerable interest in deciphering biologic differences in epithelial cells of the duct and lobules. Results of our transcriptomics study identified few major differences between these two epithelial cell types. Predominant expression of ALDH1A3 in ductal epithelial cells is interesting in the context of stem-progenitor-mature cell hierarchy of the breast ^46^. ALDH1A3 is expressed predominantly in LASP cells from which most breast cancer in BRCA1 mutation carriers is suggested to originate ^47,48^. Ductal epithelial cells also expressed higher levels of KRT14 and KRT17, which are expressed mostly in BM cells. It is possible that ducts are enriched for BM and LASP cells, whereas lobules are enriched for LHS cells and most lobular carcinomas are ER+.

Identifying age-dependent changes in breast epithelial cell gene expression has been of interest to several groups because any deviation in this process may predispose breast epithelial cells for breast tumorigenesis. Prior studies, however, utilized breast tissues from different individuals for such studies ^8,49^. Because of inter-individual differences in gene expression, results of those studies are less generalizable. Our study utilizing breast tissues from the same donor has enabled mapping of gene expression changes in ductal and lobular epithelial cells with age. PTBP1 expression is reduced at timepoint 2 compared to timepoint 1 in both cell types. PTBP1 is involved in mRNA biosynthesis and alternative splicing and its downregulation can have a major impact on cellular proteome ^30^. Similar to a previous report, basal cell marker KRT17 increased with age in ductal epithelial cells ^7^. Pathway analysis of genes commonly differentially expressed in lobular and ductal epithelial cells revealed downregulation of PKA pathway but upregulation of EIF2 pathway with aging. Studies in yeast models have shown attenuating PKA pathway extends life span and translational inhibition through phosphorylation of eIF2 increases lifespan ^50,51^. Since PKA and EIF2 pathways are also involved in breast tumorigenesis ^52,53^, whether deregulation of natural aging-dependent changes in these pathways contribute to breast tumorigenesis is an unanswered question. Collectively, data presented in this study provide an important resource generated from healthy breast tissues to derive cell type-specific chromatin accessibility and age-dependent gene expression signatures to study various diseases of the breast.

## Materials and Methods

### Normal breast tissues and genetic ancestry mapping of donors

All breast tissues from clinically breast cancer free healthy women were collected by the Komen Normal Tissue Bank (KTB) with informed consent. The study has received the approval from the institutional review board. International Ethical Guidelines for Biomedical Research Involving Human subjects were followed. KTB website describes standard operating procedure and breast biopsies were always collected from the upper outer quadrant of the breasts. Collected breast tissues were cryopreserved till use as described previously ^54^. DNA from blood of tissue donors were used for genetic ancestry mapping. Procedures for genetic ancestry mapping are described in our recent publication ^13^.

### Single cell multiome assay

Tissues from two cores of five donors were rapidly thawed and multiome assay reactions and sequencing were done in multiple batches. Nuclei were isolated from these tissues using the protocol suggested for use with the 10X Genomics chromium Next GEM Single Cell Multiome ATAC+Gene Expression protocol (CG000338). Thawed tissues were washed in PBS, minced using scalpel, and transferred to 1.5 ml microcentrifuge tubes. 300 µl of NP40 Lysis buffer was added and tissues were homogenized 15x using a Pellet Pestle (Fisher Scientific: 749625-0010). After homogenization, 1 ml of NP40 lysis buffer was added and incubated on ice for three mins. Wide bore pipet was used to mix tissues in between few times to allow better disintegration. Lysed tissue suspension was first filtered through a 70 µM strainer followed by 40 µM strainer into a 50 ml conical tube. After centrifugation for 5 mins at 500 rcf at 4 degrees, most of the supernatant was removed leaving behind 50 µl. One ml of PBS + 1% BSA + 1U/µl of RNAse inhibitor was added and kept on ice for 5 mins. After resuspension through pipet, nuclei were centrifuged at 500 rcf for 5 min at 4 degrees. After removing supernatant, nuclei were resuspend in 500 µl of PBS +1% BSA + RNase inhibitor. 5 µl of 7AAD ready-made solution (Sigma Aldrich: SML1633-1ML) was added to the above nuclei suspension. 7AAD+ nuclei were separated based on size and granularity using BD FACSMelody (or equivalent). After estimating nuclei concentration through manual counting under a microscope, next step was performed immediately as per 10X Genomics Chromium Next GEM Single Cell Multiome ATAC + Gene Expression User Guide (CG000338). ATAC and cDNA libraries were prepared using the 10X Genomics protocol (CG000338. The final ATAC and gene expression libraries were sequenced on an Illumina Novaseq 6000 sequencer, with index reads of 10 bp + 24 bp, and 100 bp paired-end reads.

### Single-cell multiome data analysis

Cell Ranger ARC v2.0 (http://support.10xgenomics.com/) was utilized to process the raw sequence data derived from the single-cell multiome libraries. Both the ATAC and gene expression FASTQ files were processed with the cellranger-arc count algorithm. The paired information of the gene expression UMIs and the count of transposition events in peaks for each barcode was used to identify the cells from the non-cell populations. The final filtered gene-cell barcode matrices and fragment files were used for further analysis with Signac ^55^ and Seurat v4 ^56–58^. Since each sample was a pool of cells from multiple subjects, souporcell ^59^ was used for genotype-free demultiplexing and the cells were assigned to their origins. Analysis with Signac and Seurat started with quality check of the cells identified. From the gene expression data, low quality cells and/or cells with extremely high or low number of detected genes/UMIs were excluded. For the ATAC-seq data, cells with low signal enrichment around transcriptional start sites, extremely high or low number of reads fallen in peaks detected were discarded. The gene expression data was normalized using SCTransform ^60^. The chromatin accessibility data was normalized applying the frequency-inversed document frequency (TF-IDF) procedure. After finishing the pre-processing and dimensionality reduction independently on the gene expression and chromatin accessibility data, the closest neighbors of each cell in the data were calculated based on a weighted combination of gene expression and chromatin accessibility similarities. The weighted nearest neighbor graph (WNN) calculated were used for cell clustering and visualization.

To annotate each cell population from the analysis, automatic annotation using SingleR together with manual annotation with known marker genes were employed. CoveragePlot function from Signac was used to plot chromatin accessibility for specific genomic regions.

### Single-nuclei data analysis

snRNA-seq data was first processed using CellRanger 7.0.1 (http://support.10xgenomics.com/). The feature-cell barcode matrices generated from CellRanger was used for further analysis with the R package Seurat v4 ^56–58^. The integrated single-cell multiome data was used as a reference to annotate the snRNA-seq data. FeaturePlot_scCustom function in scCustomize ^61^ was used to generate the gene expression plots.

### Spatial Transcriptomics analyses

FFPE sections from donors with two time tissue donations were selected for the study. Each slide contained two 5 micrometer sections from FFPE blocks. Each slide represented one donor with 2 barcodes representing 2-time donations. Each donor had three repeats (3 slides per donor). The sections were cut with Leica DB80 LS blades (Leica #14035843488) on a rotary microtome instrument (Leica RM2125 RTS) and placed on the center of a Superfrost Plus Microscope slide (Fisher scientific #1255015). Tissue sections were placed in the center of the slide and be no larger than 35.3 mm x 14.1 mm.

Regions of interest (ROIs) were selected after staining the slides with pan-keratin, alpha-SMA and FABP4 antibodies. All ROIs passed a sequencing quality control assessment. Next, negative control probes were used to estimate background and downstream gene detection and to remove outliers. The limit of quantification (LOQ) of each ROI was calculated using the geometric mean and geometric standard deviation of the negative control probes to identify genes detected above background in the experiment. All ROIs passed LOQ-based filtering with more than 1% of genes detected. Gene filtering was also performed, resulting in 10,270 remaining targets that were detected above the LOQ in 10% or more ROIs. 10,270 genes remained for data analysis from 48 ROIs. Upper quartile (Q3) normalization was performed for genes in each segment. Quality control and normalization was performed using GeoMxTools v3.0.1.

### Statistical analyses of Spatial transcriptomics data

Dimension reduction analysis was performed in R v4.2.1 using the following packages: ‘FactoMineR’ v2.6, ‘Rtsne’ v0.16, and ‘UMAP’ v0.2.9.0. Differential gene expression analysis was performed on a per-gene basis, modeling log-transformed, normalized gene expression using either a linear mixed-effect model (LMM) for study-wide comparisons or a linear model for donor-specific comparisons with GeoMxTools. LMMs are used to account for the sampling of multiple ROI/AOI segments per tissue and non-independence of the data. For the study-wide pairwise comparisons between the ducts vs lobules, the following LMM was used: gene ∼ ROIType + (1|Tissue). For comparing ducts vs lobules within the same donor tissues, the following linear model was used: gene ∼ ROIType. A false discovery rate (FDR) correction was applied to p-values. To avoid inter-sample variability impacting data interpretation, the following seven analyses were performed with the first four analyses restricted to sample number 3. Question 1: What are the differences between the duct and gland at the first timepoint?; Question 2: What are the differences between the duct and gland at the second timepoint?; Question 3: What are the differences in the duct between timepoints 1 and 2?; Question 4: What are the differences in the gland between timepoints 1 and 2?; Question 5: What are the differences between ducts and glands across all donors?; Question 6: What are the differences between ducts and glands at timepoint 1?; Question 7: What are the differences between ducts and glands at timepoint 2?

Spatial deconvolution was performed using the SpatialDecon package in R, v1.6.0. Spatial deconvolution requires the use of a cell profile matrix derived from scRNA-seq. For this analysis, we used gene signatures derived from multiome data Tables S4-S9. Differential abundance analysis was performed on the results of spatial deconvolution using the same approach as differential gene expression.

Pathway analysis was performed using the GSVA v1.44.5 R package with the KEGG Brite database. 796 gene sets were scored, where each gene set contained between 5 and 500 genes. Differential gene set enrichment analysis was performed on the results using the same approach as differential gene expression. Ingenuity pathway analyses was used to determine pathways altered due to aging.

## ACKNOWLEDGMENTS

We thank the countless number of women who donated normal and malignant breast tissues for research. We also thank the volunteers who facilitated this tissue collection. Special thanks to members of the Komen Tissue Bank including Ms. Jill Henry, Alison Hughes, Pam Rockey, and Julia Rose von Arx, as well as the IU Simon Cancer Center tissue procurement facility for providing tissues and related data. We also thank Dr. David Scoville of Nanostring technology for processing GeoMx data.

## AUTHOR CONTRIBUTIONS

Conception and design: HN

Development of methodology: PBN, DC, ASK, GJ, PCM, HG, CE, RG, FN, YL, HN

Acquisition of data: PBN, FN, HG

Analysis and interpretation of data: PBN, AKS, CE, GJ, FN, HG, YL, AMS, HN

Writing, review and/or revision of the manuscript: PBN, DC, RG, HG, FN, AMS, HN

Administrative, technical, or material support: AMS, YL, HN

Study supervision: HN, YL

## Data availability

The majority of data are included in the manuscript. High throughput data are available through NCBI database with SuperSeries accession number GSE244594. In addition, these data are publicly available through CellXGene database of Chan-Zuckerberg Initiative.

