## Supplementary figures for "Single nuclei chromatin accessibility and transcriptomic map of breast tissues of women of diverse genetic ancestry"

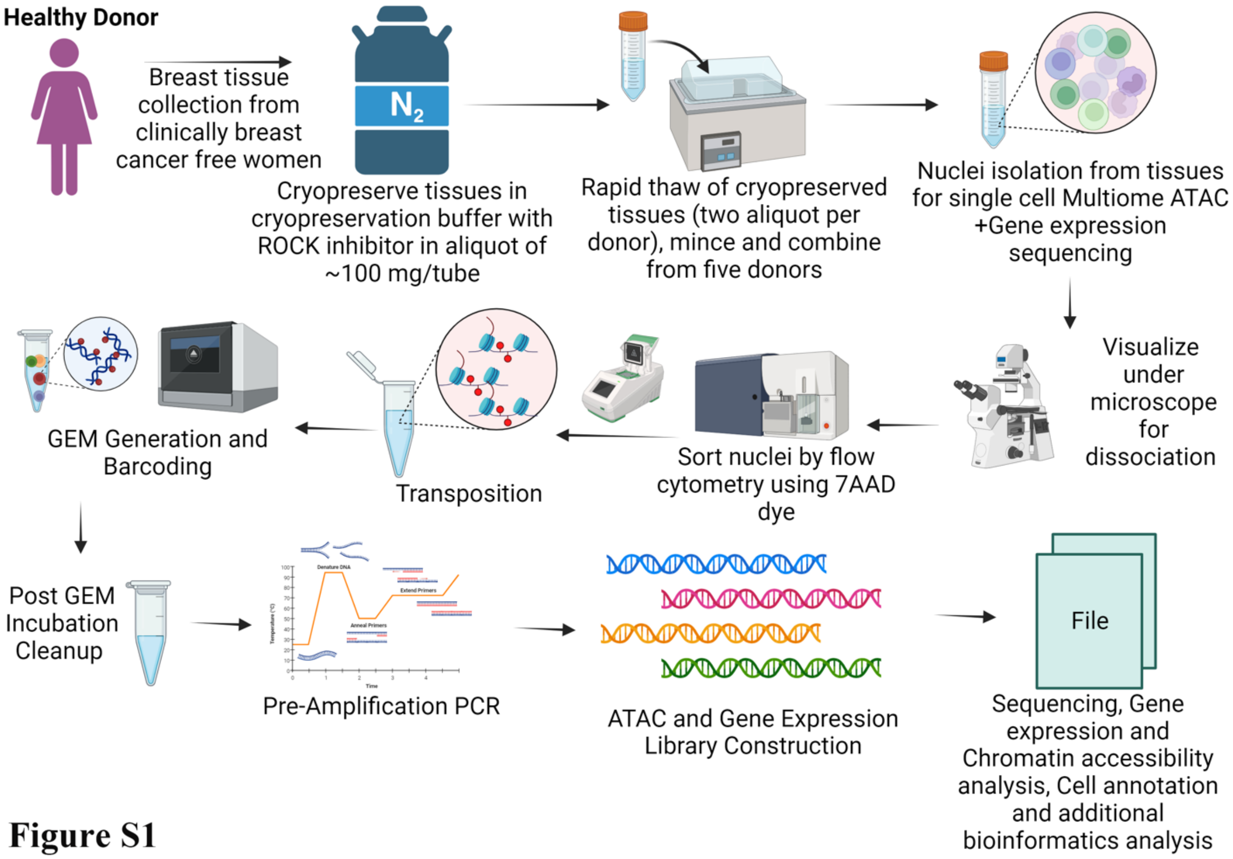


**Figure S1**

**Figure S1:** Experimental design to generate snATAC-seq and snRNA-seq data.

**a**

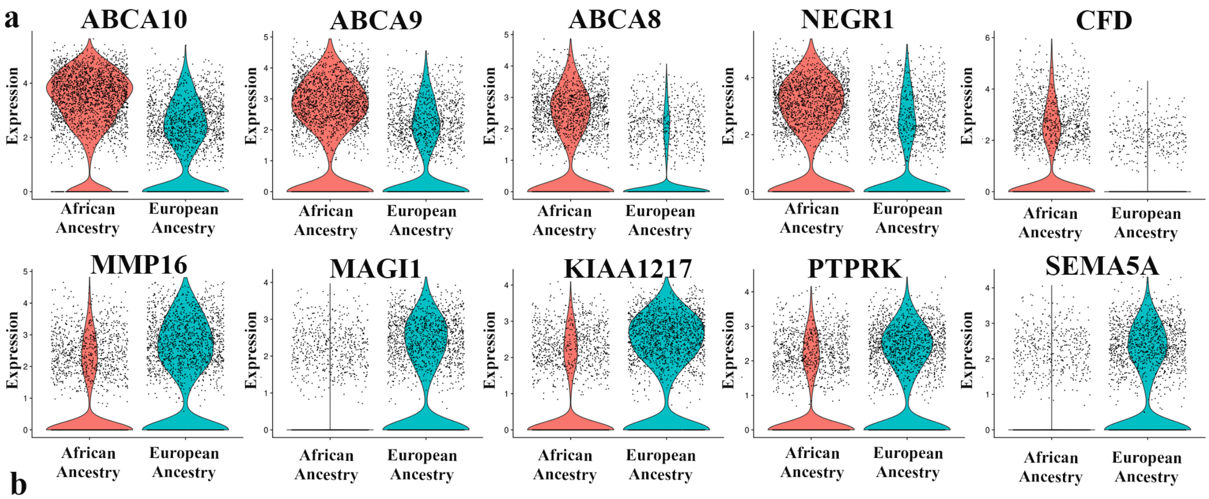


**b**

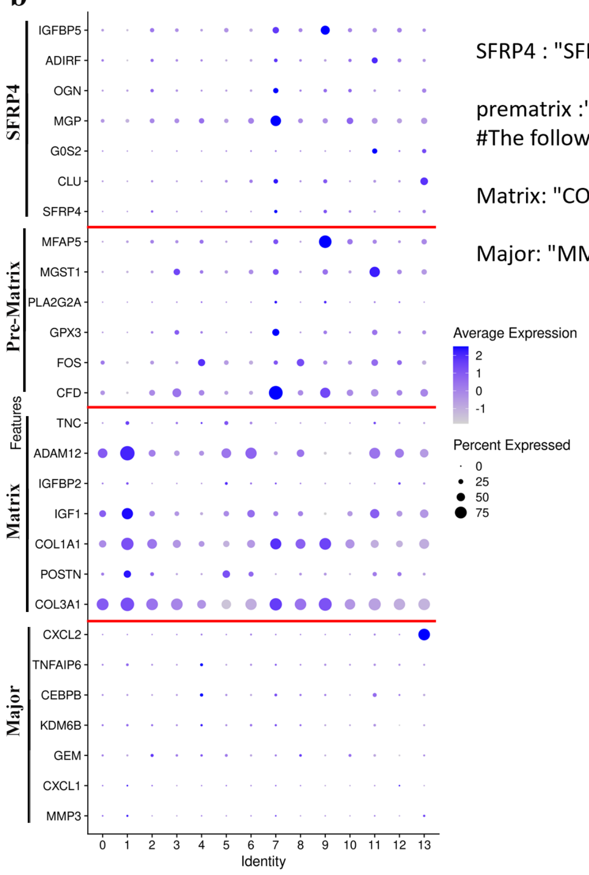


SFRP4 : "SFRP4","CLU","G0S2","MGP","OGN","ADIRF","IGFBP5"

prematrix :"CFD","FOS","GPX3","PLA2G2A","MGST1","MFAP5"
#The following requested variables were not found: WISP2

Matrix: "COL3A1","POSTN","COL1A1","IGF1","IGFBP2","ADAM12","TNC"

Major: "MMP3","CXCL1","GEM","KDM6B","CEBPB","TNFAIP6","CXCL2"

**Figure S2**

**Figure S2:** Differences in expression of fibroblast-enriched genes in breast tissue fibroblasts of African ancestry compared to European ancestry. Expression levels of genes that classify fibroblasts into four subtypes are also shown.

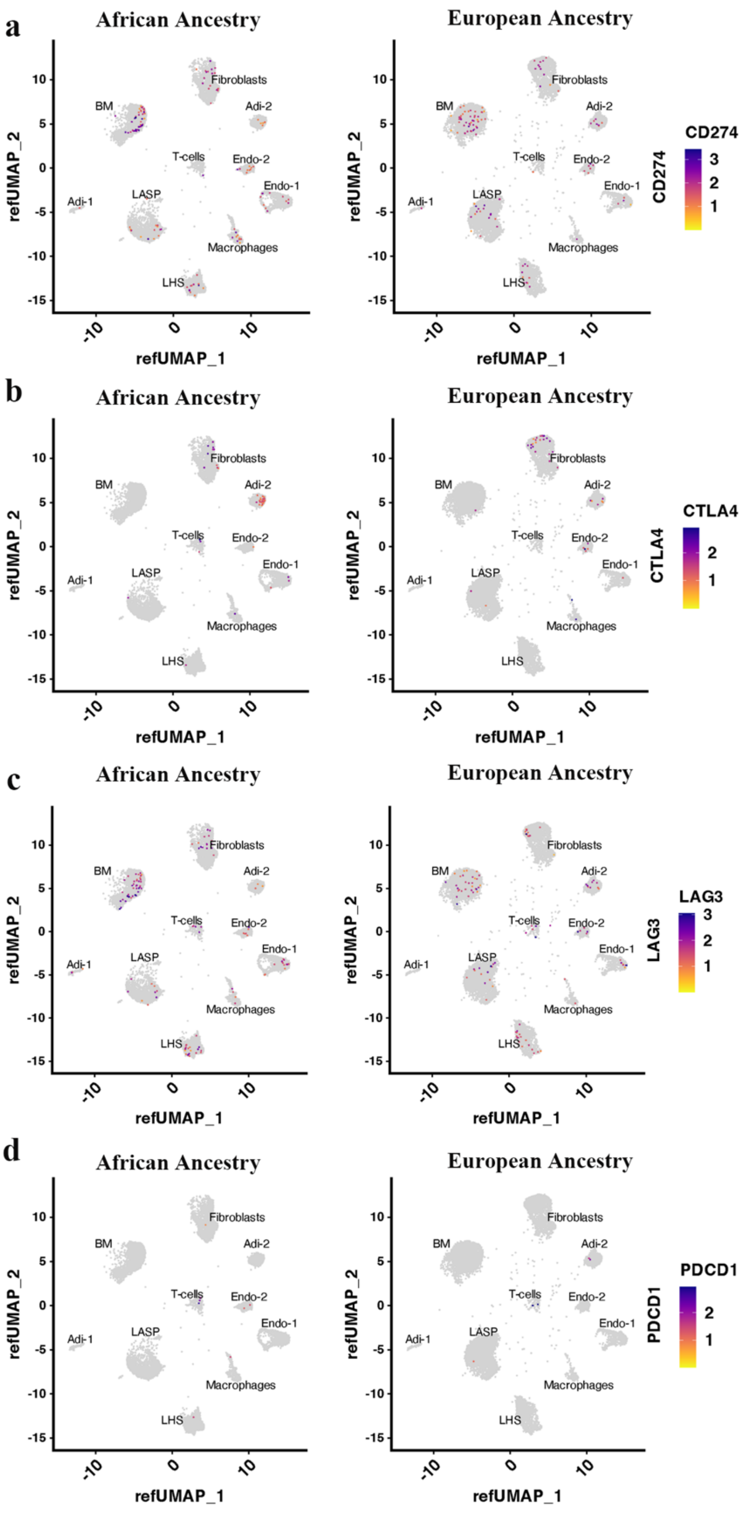
a
African Ancestry
European Ancestry
CD274
b
African Ancestry
European Ancestry
CTLA4
c
African Ancestry
European Ancestry
LAG3
d
African Ancestry
European Ancestry
PDCD1
refUMAP_1
refUMAP_2
Fibroblasts
BM
Adi-2
T-cells
Endo-2
Endo-1
Adi-1
LASP
Macrophages
LHS

Figure S3

**Figure S3:** Expression pattern of exhausted T cell markers in breast tissues of women of African and European ancestry

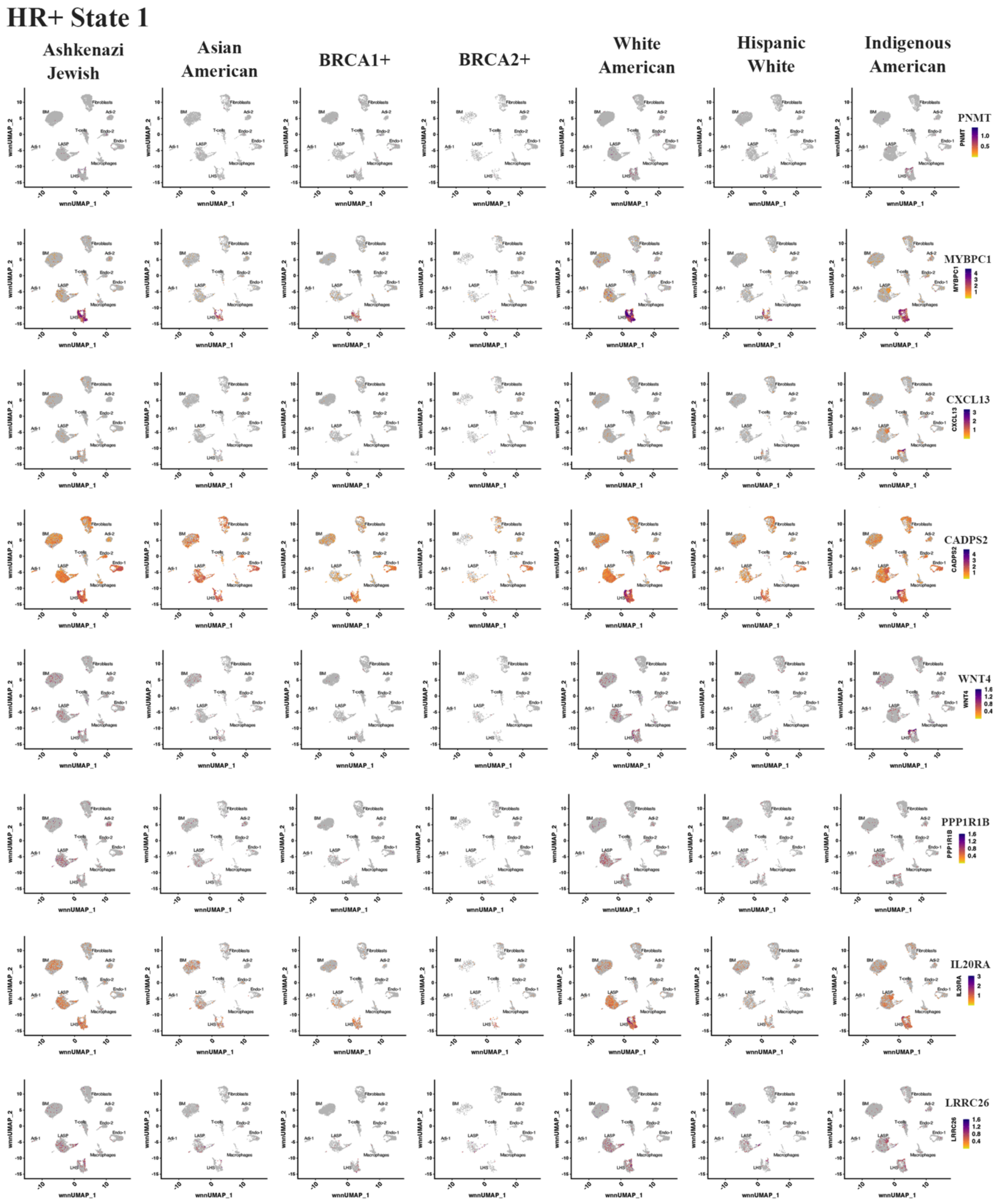
HR+ State 1
Ashkenazi Jewish
Asian American
BRCA1+
BRCA2+
White American
Hispanic White
Indigenous American
PNMT
MYBPC1
CXCL13
CADPS2
WNT4
PPP1R1B
IL20RA
LRRC26

Figure S4a

### HR+ State 2

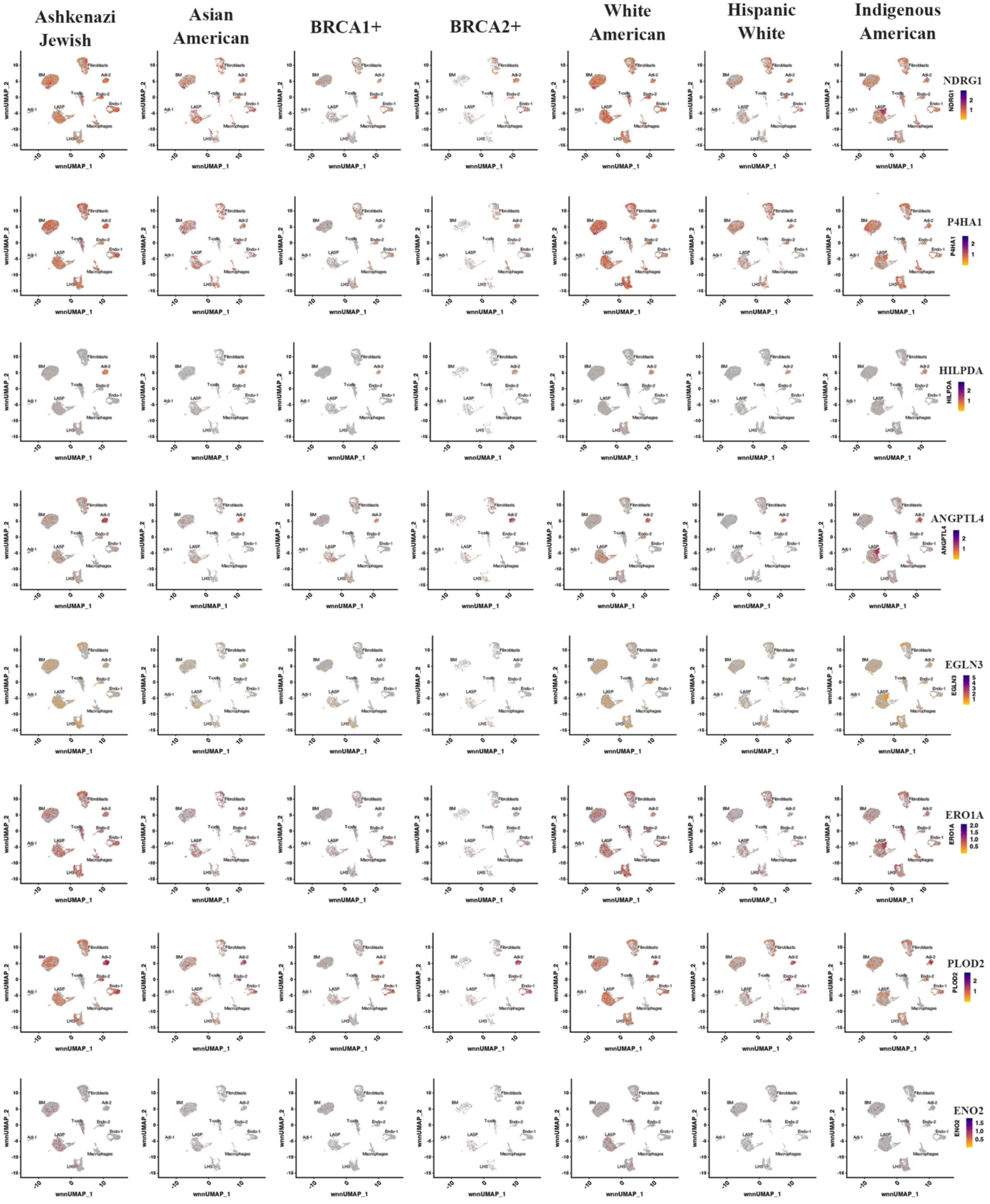


**Figure S4b**

**Figure S4:** Expression pattern of markers suggested to classify hormone responsive (HR) cells into ER/PR-dependent HR State-1 (a) and hypoxia/pro-angiogenic HR State-2 (b).

a

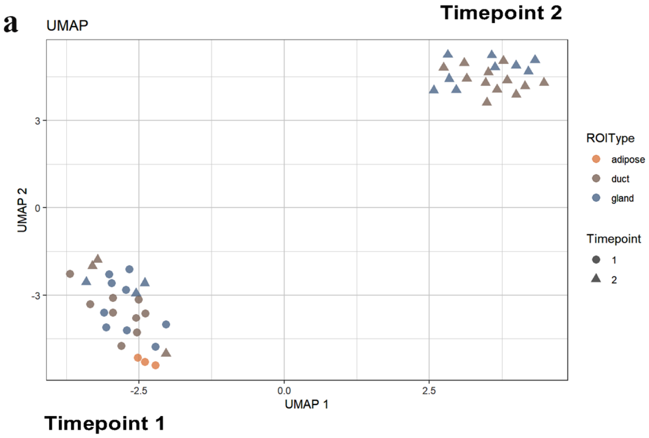


### Samples used

| Sample | Time 1 age (BMI) | Time 3 age (BMI) |
|---|---|---|
| 1 | 34 (31.7) | 44 (32.2) |
| 2 | 60 (29.3) | 70 (27.8) |
| 3 | 27 (23.4) | 37 (33.5) |

b

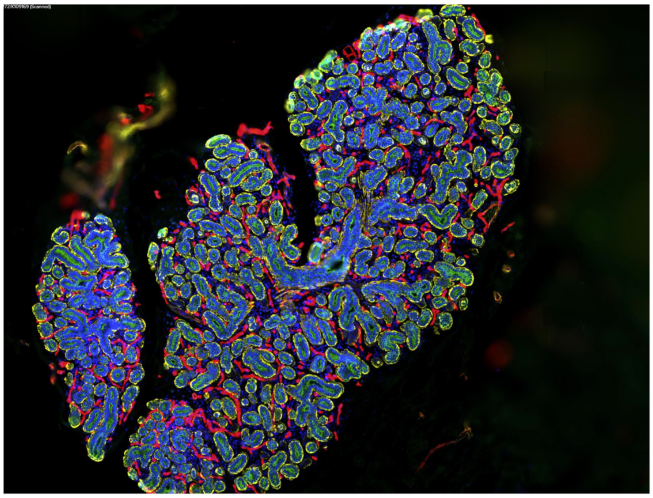

Green –pan-keratin- epithelial cells
Red- FABP4- adipocytes
Yellow- Smooth muscle actin

Five pairs of samples collected 10 years apart
Three pairs have provided data.

c

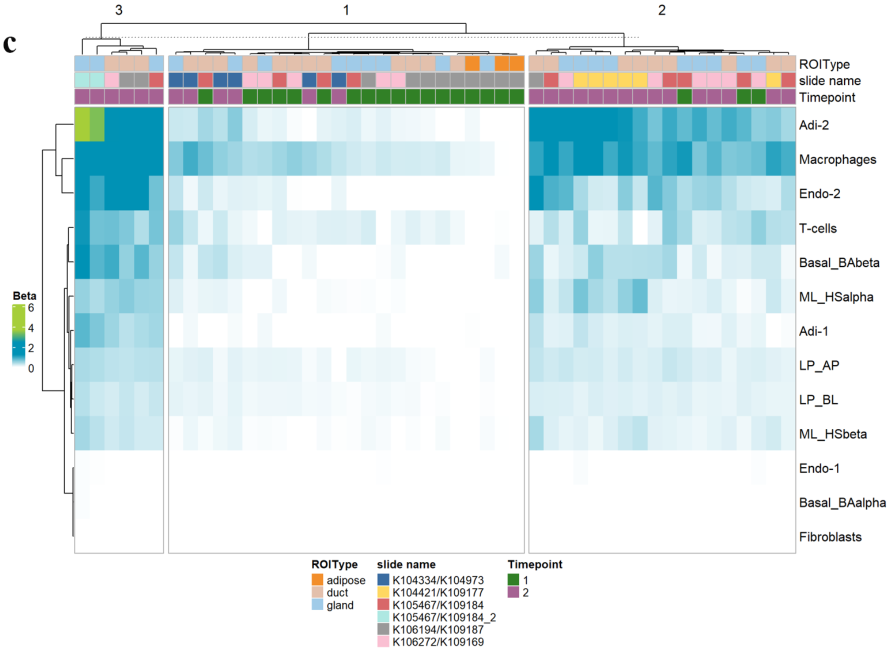


**Figure S5**

**Figure S5:** a) UMAP showing differences in gene expression patterns between timepoint 1 and timepoint 2. Age and BMI of donors at two timepoints of tissues collected for spatial transcriptomics are also indicated. b) Staining pattern of breast tissues with antibodies against pan-keratin, FABP4 and smooth muscle actin. c) Deconvolution of spatial transcriptomics data show elevated Adi-2, macrophages and Endo-2 at timepoint 2 compared to timepoint 1 in most samples.

### Duct Vs Lobule: Timepoint 1

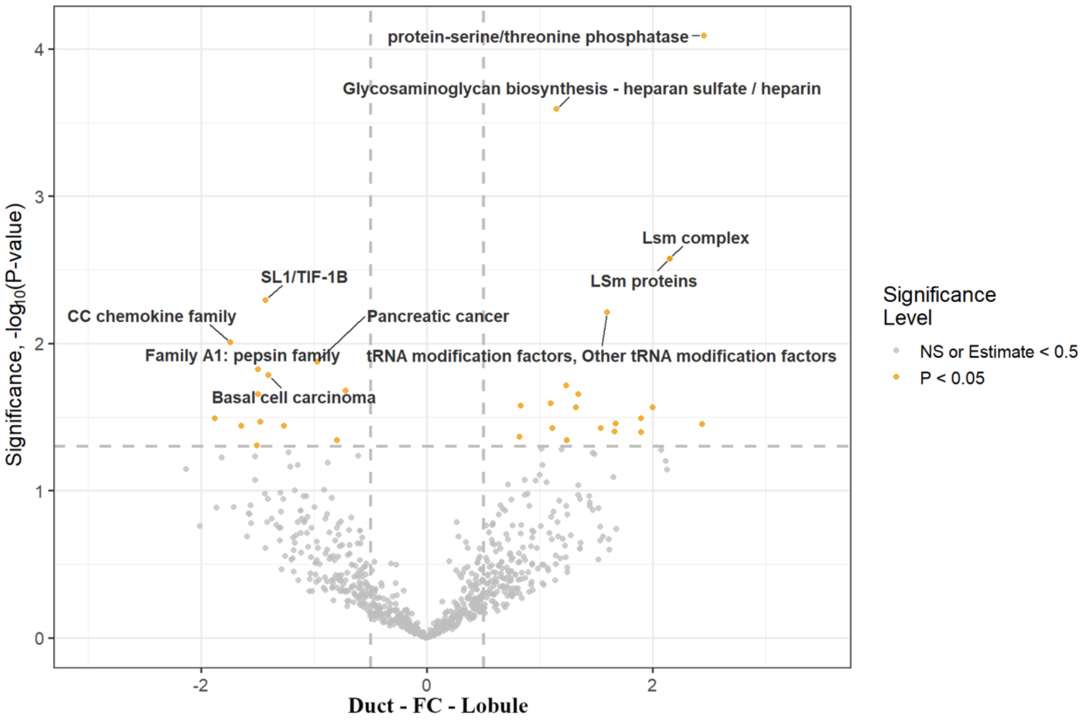


### Duct Vs Lobule: Timepoint 2

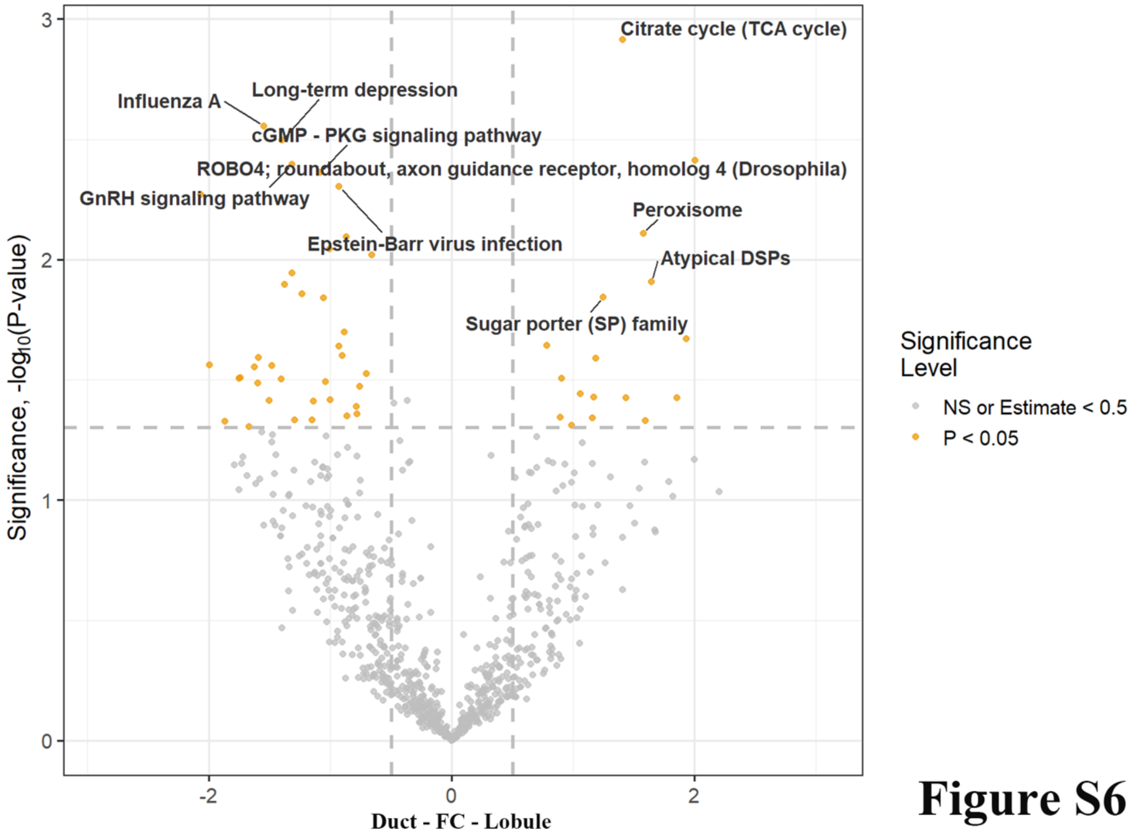


**Figure S6**

**Figure S6:** Differences in signaling pathways in ductal and lobular epithelial cells.

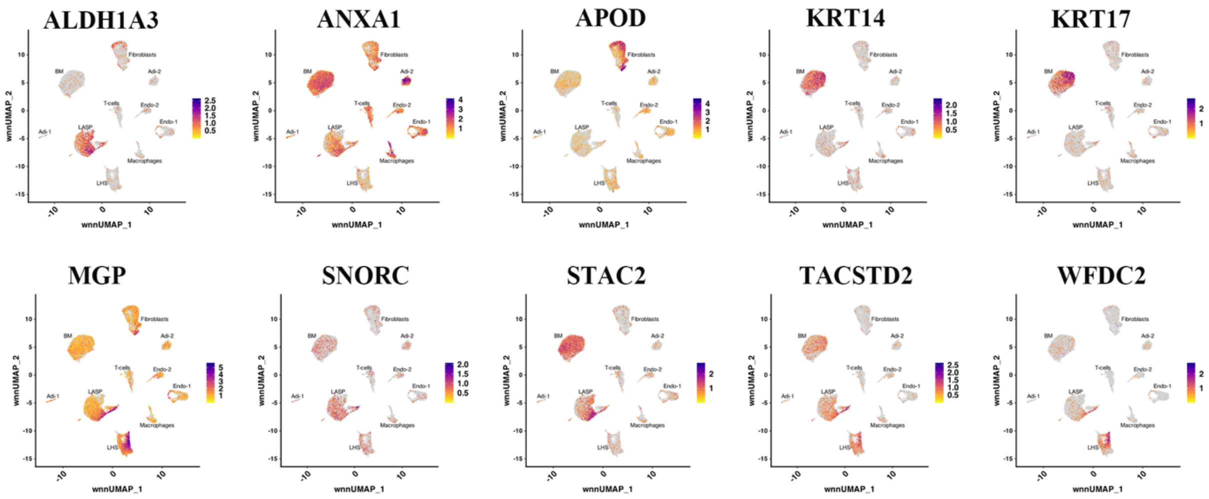
ALDH1A3
ANXA1
APOD
KRT14
KRT17
MGP
SNORC
STAC2
TACSTD2
WFDC2
Fibroblasts
BM
Adi-2
T-cells
Endo-2
Endo-1
Adi-1
LASP
Macrophages
LHS
wnnUMAP_1
wnnUMAP_2

Figure S7

**Figure S7:** Expression pattern of 10 genes that showed differential expression in ductal epithelial cells compared to lobular epithelial cells assessed using multiome data.

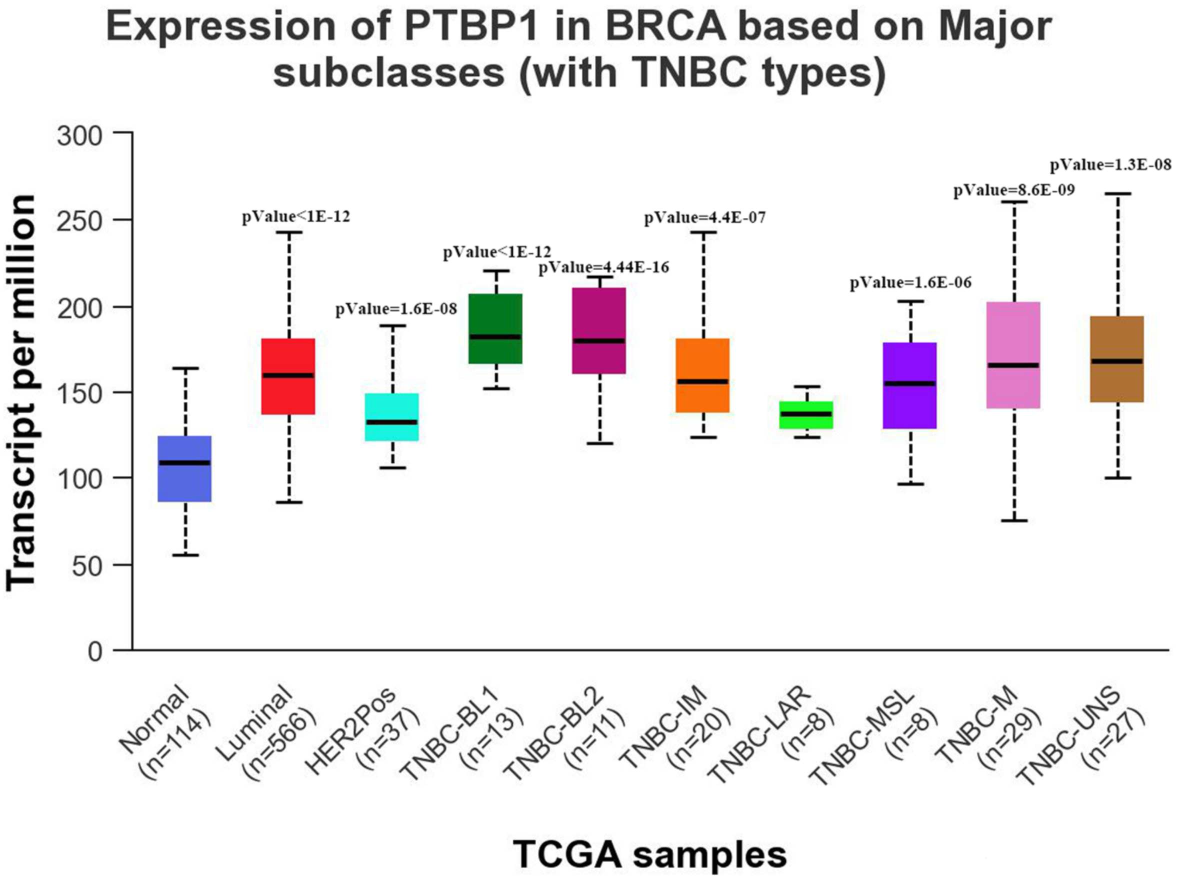
Expression of PTBP1 in BRCA based on Major subclasses (with TNBC types)
Transcript per million
300
250
200
150
100
50
0
pValue<1E-12
pValue=1.6E-08
pValue<1E-12
pValue=4.44E-16
pValue=4.4E-07
pValue=1.6E-06
pValue=8.6E-09
pValue=1.3E-08
Normal (n=114)
Luminal (n=566)
HER2Pos (n=37)
TNBC-BL1 (n=13)
TNBC-BL2 (n=11)
TNBC-IM (n=20)
TNBC-LAR (n=8)
TNBC-MSL (n=8)
TNBC-M (n=29)
TNBC-UNS (n=27)
TCGA samples

**Figure S8**

**Figure S8:** PTBP1 whose expression in normal breast epithelial cells was reduced in timepoint 2 compared to timepoint 1, is overexpressed in all breast cancer subtypes compared to normal breast.
