## Supplementary material for "Single nuclei chromatin accessibility and transcriptomic map of breast tissues of women of diverse genetic ancestry": Table S2

**Table S2: Number of donor tissues, nuclei and transcripts per nuclei in each group.**

| Genetic ancestry/mutation | Number of samples | Number of nuclei/cell | Number of gene per nuclei/cell |
|---|---|---|---|
| Ashkenazi | 22 | 13,959 | 1246 |
| Asian | 10 | 3,178 | 1,689 |
| European-non-Ashkenazi | 20 | 13,269 | 1,162 |
| Hispanic-White | 10 | 4,116 | 1,112 |
| Indigenous American | 10 | 10,575 | 1,251 |
| BRCA1 | 6 | 4695 | 735 |
| BRCA2 | 5 | 1575 | 653 |
| African American* | 10 | 17444 | 1215 (median) |
| European* | 10 | 19194 | 1257 (median) |

* snRNA-seq only
